# Non-lytic replicating viral delivery of an IL15 superagonist enhances antitumor immunity and extends survival in glioblastoma

**DOI:** 10.1101/2024.10.14.618095

**Authors:** Alexander F. Haddad, Atul Saha, Jordan Spatz, Sara A. Collins, Isabella Lovalvo, Sabraj Gill, Megan L. Montoya, Poojan Shukla, Jinpyo Hong, Elaina Wang, Pavlina Chuntova, Meeki Lad, Robert Osorio, Jia-Shu Chen, Saket Jain, Eric Chalif, Noriyuki Kasahara, Manish K. Aghi

## Abstract

Glioblastoma (GBM) is the most lethal primary brain neoplasm due to its highly immunosuppressive microenvironment and resistance to conventional therapies. To overcome this challenge, we engineered a replicating retrovirus (RRV) to deliver a superagonist interleukin-15 receptor-linked fusion protein (RLI) directly to tumor cells, engineering them into local immunotherapy biofactories. This strategy leverages the tumor-selective replication of RRV to achieve localized and sustained RLI expression within the tumor microenvironment. In two orthotopic poorly immunogenic GBM mouse models, intratumoral administration of RRV RLI significantly reduced tumor growth and prolonged survival compared to controls, with some mice achieving long-term remission and demonstrating immunologic memory upon rechallenge. Transcriptomic and flow cytometric analyses revealed that RRV RLI treatment enhanced infiltration and activation of CD8⁺ T cells, NK cells, and upregulated antigen presentation pathways within the tumor microenvironment. Depletion studies indicated that the therapeutic efficacy of RRV RLI is dependent on both CD4⁺ and CD8⁺ T cells. Notably, combining RRV RLI with the GBM standard of care chemotherapeutic agent temozolomide (TMZ) synergistically improved survival outcomes. Subsequent single-cell RNA and T cell receptor sequencing identified enhanced effector cell activation, antigen presentation, and clonal T cell expansion in the combination therapy group. Further T cell receptor analysis and clustering implied a tumor-specific immune response rather than one targeting the viral delivery vehicle, suggesting that this therapeutic approach could be reapplied without eliciting anti-vector immunity. Our findings suggest that RRV-mediated delivery of RLI effectively transforms GBM tumors into immunostimulatory hubs, eliciting a potent anti-tumor immune response. This novel viral immunotherapy holds significant promise for clinical translation in the treatment of GBM and other difficult-to-treat solid tumors.

## Introduction

Glioblastoma (GBM) is a devastating brain cancer and carries a poor prognosis despite standard-of-care treatment, including surgical resection, radiation, and chemotherapy.^1^ The median survival for newly diagnosed GBM is 15 months, with only modest improvements over the past decade and a need for novel therapeutics.^2^ ^3–5^ Despite success in other cancer types, systemically delivered single-agent immunotherapy has thus far seen limited success in treating GBM,^4^ due to the tumor’s uniquely immunosuppressive microenvironment.^3,4,6–9^ Several characteristics contribute to the immunodeficient and immunosuppressive tumor microenvironment (TME) of GBM, including the downregulation of MHC I,^7–9^ low tumor mutational burden (TMB),^10^ tumor intrinsic signaling pathways,^11^ and infiltration of immunosuppressive myeloid cells.^12^ A key contributor to the immunosuppression seen in GBM is the low number of tumor-infiltrating T cells, which are often exhausted and dysfunctional.^6,13,14^ GBM patients also have low systemic T cells due to their sequestration in the bone marrow, compounding the low anti-tumor T cell activity seen.^15^

Systemic immunotherapy for GBM faces additional challenges. The doses needed to overcome the severe immunosuppression^16^ and cross the blood-brain barrier may be high enough to cause systemic toxicities such as autoimmune side effects or cytokine storm.^16–18^ Consequently, there is a growing interest in local immunotherapies that deliver immunostimulatory proteins like cytokines directly to tumor cells.^19,20^ While local immunotherapies investigated to date have taken many forms, such as nanoparticles, scaffolds, and hydrogels, one of the most promising is the use of viral vectors to deliver therapeutic transgenes, such as a suicide gene or local immunotherapy directly to the tumor microenvironment.^19,20^

Replicating retrovirus (RRV) is a promising viral vector for cancer gene therapy that selectively infects, stably integrates, and replicates in tumor cells.^21^ Unlike other viral vectors, RRV is relatively immunologically silent, allowing widespread tumor dissemination with minimal immune clearance.^22^ The RRV Toca 511 (vocimagene amiretrorepvec), which delivers the suicide gene yeast cytosine deaminase (CD), has been used in multiple glioblastoma clinical trials, including a randomized multicenter phase III clinical trial, which despite failing to improve survival compared to standard of care for recurrent GBM, exhibited an excellent safety profile.^23,24^ Molecular profiling of treated patients further emphasizes the tumor-selectivity and safety of the RRV backbone, highlighting its promise as a highly clinically translatable vector to deliver a range of transgenes.^23^

Immunostimulatory cytokines, such as interleukin-2 (IL2) and interleukin-15 (IL15), represent promising candidates for transgene therapy due to their capacity to enhance the proliferation of Natural Killer (NK) and T cells.^25^ Of particular interest is IL15, which exhibits a favorable profile by stimulating the expansion of NK and T cells while also promoting their cytotoxic function and release of additional inflammatory cytokines.^25^ Unlike IL-2, IL15 does not stimulate T regulatory (Treg) cells which can suppress anti-tumor immunity, thereby offering a therapeutic advantage.^25^ Clinical utility of IL15 alone is in part limited by its complex signaling pathway, involving trans-presentation, a novel mechanism of delivery unique to IL15 as IL15 primarily exists bound to the high-affinity IL15Rα leading to IL15/IL15Rα complexes that are shuttled to the cell surface of NK- and T cells, where they can stimulate opposing cells such as myeloid cells through the β/γC receptor complex. In oncology patients, the expression of IL15Rα is frequently downregulated, and IL15 has a relatively short half-life, further complicating its application as a therapeutic agent.

These limitations have inspired the development of superagonists which can directly bind to effector cells and have increased stability.^26–28^ One such superagonist is receptor-linked–IL15 (RLI), which is bound to the sushi domain of the IL15 receptor alpha, allowing it to bind directly to IL15 receptors on NK and T cells and improving stability. Systemically administered RLI has been shown to slow tumor growth and promote anti-tumor immune activity in multiple preclinical cancer models.^26–28^

Given the promising preclinical results associated with systemic RLI treatment, we hypothesized that local RLI therapy delivered via RRV would stimulate an anti-tumor immune response. In the present study, we successfully engineered an RRV expressing RLI and demonstrated its ability to increase survival in poorly immunogenic mouse models of GBM in a T cell-dependent manner. RRV RLI also synergized with GBM standard-of-care chemotherapy with leukocyte single-cell sequencing revealing clonal T cell expansion, enhanced antigen-driven immune response, and favorable shifts in myeloid populations when the two treatments were combined. This approach provides a novel viral cancer immunotherapy for GBM with a high potential for clinical translation.

## Results

### IL15 expression is associated with immune signaling and T cell Infiltration in glioblastoma, with implications for gene therapy

To better understand the applicability of IL15 as a cancer immunotherapy, we examined baseline IL15 expression across various cancer types and investigated differences between IL15 high and low glioblastoma (GBM) tumors. Analysis of bulk RNA sequencing data revealed that gliomas of all grades, including GBM, exhibited among the lowest levels of IL15 expression across cancer types, whereas thyroid cancer showed the highest expression (**Fig. 1A**). In GBM tumors with high IL15 expression, we observed significantly increased expression of immune and inflammatory genes such as *IL2RA, CD70, IL6, and CCL11*, as well as immunoglobulin genes *IGKV1D-17* and *IGHV3-72* (**Fig. 1B**). Subsequent gene ontology analysis indicated upregulation of pathways associated with immunoglobulin production, phagocytosis, antigen binding, receptor-ligand activity, and molecular mediators of immune response (**Fig. 1C**).

**Fig 1.**
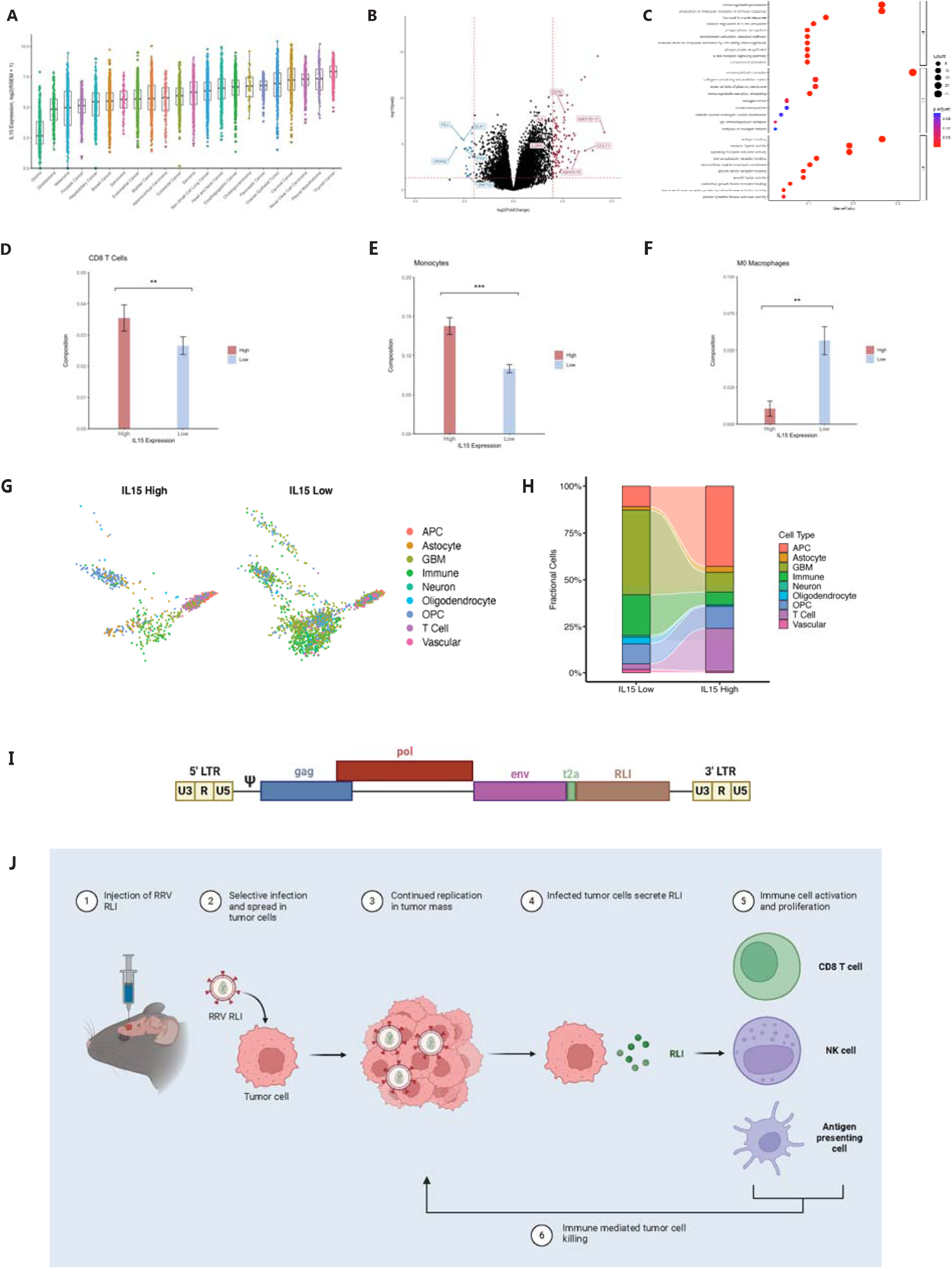
Human IL15 expression in various cancers and its biologic effects in GBM informs the decision to create RRV RLI. (**A**) IL15 expression across cancer types. (**B**) Volcano plot demonstrating differential gene expression between IL15 high and low glioblastoma tumors. (**C**) Gene ontology analysis comparing IL15 high and low glioblastoma tumors. **D**) Calculated CD8^+^ T cell infiltration based on bulk RNA sequencing with increased infiltration in IL15 high samples. (**E**) Calculated monocyte infiltration based on bulk RNA sequencing demonstrating increased infiltration in IL15 high tumors. (**F**) Calculated M0 macrophage infiltration based on bulk RNA sequencing showing decreased infiltration in IL15 high tumors. (**G**) Single cell RNA sequencing UMAP of human glioblastoma tumors comparing IL15 high vs. low tumors. (**H**) Alluvial plot demonstrating changes in populations between human IL15 high and low samples with increased T cell and APC infiltration in an IL15 high state. (**I**) Schematic of the RRV RLI construct with RLI placed after the RRV env protein. (**J**) Proposed mechanism of RRV RLI treatment.

CIBERSORT analysis of 22 different immune cell populations revealed a significant increase in infiltrating CD8⁺ T cells and monocytes in the IL-15 high tumors, along with a decrease in infiltrating M0 macrophages (**Fig. 1D–F**). Analysis of available single cell RNA sequencing data from four resected GBM tumors supported these findings, demonstrating increased infiltration of T cells and antigen-presenting cells (**Fig. 1G and H**).^29^ Together, this data identified GBM as a IL15-deficient tumor with a more favorable anti-tumoral immune microenvironment in those GBMs that happened to have high IL15 expression, implicating IL15 as a promising gene therapy strategy in GBM.

As a result, we generated RRV RLI, an RRV expressing an optimized RLI insert following a T2A cleavage site on the RRV backbone as previously described in the literature (**Fig. 1I**).^21^ RRVs replicate well in tumors due to their reliance on two hallmarks of cancer, sustained proliferative signaling and immunosuppression in the tumor microenvironment (**fig. S1A**).^30^ As such, we utilized RRV RLI to convert tumor cells into biofactories for secreted RLI, aiming to enhance the activation and proliferation of CD8^+^ T cells, NK cells, and antigen-presenting cells and generate an anti-tumor immune response (**Fig. 1J**).

### RRV RLI efficiently infects and spreads in murine GBM, driving the secretion of functional RLI

^2130^We first sought to understand viral infectivity kinetics in mouse orthotopic GBM tumors *in vivo* through direct intratumoral injection. We confirmed that SB28 tumors can support the replication of an RRV expressing the emerald (EMD) protein *in vivo*, with on average 85.5% of intracranial SB28 GBM cells expressing the EMD protein at tumor endpoint after direct intracranial injection, similar to a 4% ex vivo-transduced tumor (**Fig. 2A**, example gating **Fig. 2B**). This also confirmed the efficacy of intracranial injection for viral delivery. We then sought to assess viral replication and stability of the RRV RLI viral construct in cultured murine glioblastoma tumor cell lines. **Fig. 2C** demonstrates the ability of RRV RLI to efficiently spread through cultured SB28 cells beginning with multiplicities of infection (MOIs) of 1 and 0.1. Transduced cells exposed to azidothymidine (AZT), which inhibits viral spread, showed no increase in RRV RLI over time, as expected (**Fig. 2C**). Similar kinetics of RRV RLI spread were noted in cultured Tu-2449 murine GBM cells (**Fig. 2D**). Given the potential for insert dropout and loss, the stability of RRV RLI was confirmed using polymerase chain reaction (PCR) across the RLI insert site (**fig. S1B**) every two weeks for eight weeks of total culture, with no evidence of RLI insert loss (**Fig. 2E**).

**Fig 2.**
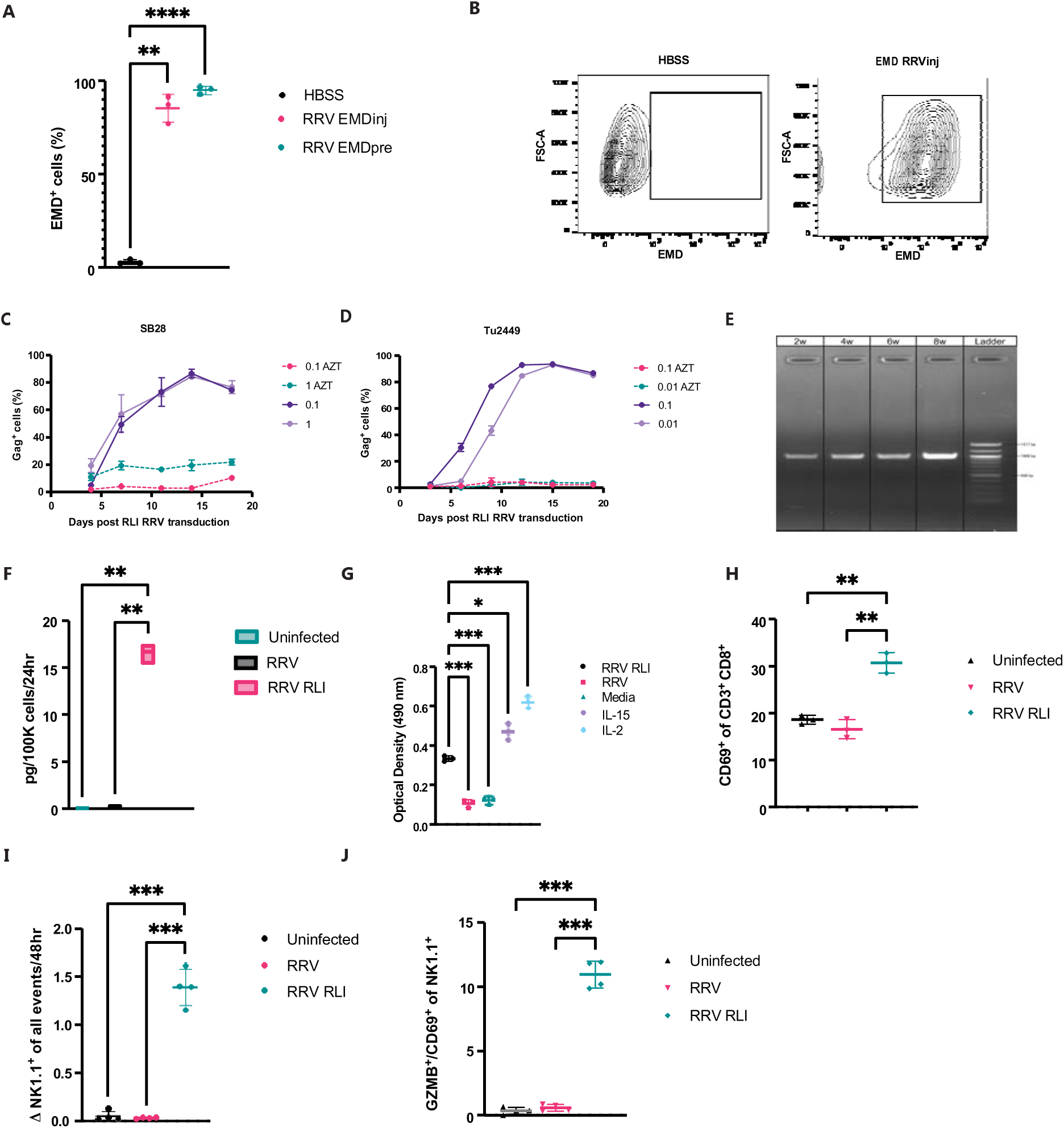
Validation of the biologic effects of RRV RLI in cultured glioblastoma cells. (**A**) *in vivo* spread of RRV-EMD in the SB28 GBM murine tumor model at endpoint. There was no significant difference between RRV-EMD injection (RRV-EMDinj) and RRV-EMD ex vivo pre-transduction (RRV-EMDpre) (85.5% vs. 95.3% EMD+ cells, p=0.14, Welch’s t-test). Both RRV-EMDinj (p<0.01) and RRV-EMDpre (p<0.001) demonstrated significantly higher infection rates compared to HBSS injection control. (**B**) Example gating of RRV EMD *in vivo* spread. (**C**) *in vitro* RRV RLI replication in SB28 is efficient, reaching >80% tumor cell infection by day 15 regardless of starting multiplicity of infection (MOI) as measured by the expression of viral Gag protein on flow cytometry with no change in AZT control groups. (**D**) *in vitro* RRV RLI replication in Tu2449 is similarly effective, reaching >90% tumor cell infection by day 15 regardless of starting multiplicity of infection (MOI) as measured by the expression of viral Gag protein on flow cytometry with no change in AZT control groups. (**E**) RRV RLI demonstrates stability and lack of transgene dropout after 8 weeks in culture in the SB28 cell line (**fig. S1B**). (**F**) RRV RLI stimulates production of 16.72 pg/100K cells over 24 hours in the SB28 cell line with minimal production in RRV (no transgene, 0.2 pg/100K cells/24hr)) or uninfected control cells (0.02 pg/100K cells/24hr). (**G**) RLI produced by SB28 cells is functional and stimulates the growth of cytokine dependent CTLL-2 cells as compared to RRV (p<0.001, Welch’s t-test) and media control (p<0.001, Welch’s t-test). Nanomolar matched (0.361 nm) recombinant IL15 (p=0.01, Welch’s t-test) and IL-2 (p<0.001, Welch’s t-test) were used as positive controls. (**H**) Co-culture of CD8 T cells with infected SB28 tumor cells demonstrates increased CD69 after 48 hours compared to RRV and uninfected cells (p<0.01, Welch’s t-test). (**I**) Co-culture of NK cells with infected SB28 tumor cells demonstrates increased change in NK cell frequency relative to all events at 48 hours when compared to RRV infected and uninfected (1.4 vs. 0.05, vs. 0.03, p<0.001, Welch’s t-test). (**J**) In NK cell: SB28 tumor cell co-culture, there was increased double positivity for GZMB and CD69 at 24 hours (p<.005, Welch’s t-test).

After confirming viral kinetics and stability, we aimed to investigate the production and function of RLI produced by infected tumor biofactories. Using an IL15 ELISA, we measured that SB28 cells infected with RRV RLI secreted RLI at an average rate of 16.7 pg/cell/24hours (**Fig. 2F**). To assess whether the IL15 secreted by RRV RLI-transduced murine GBM cells retained its canonical functions, we performed a series of functional assays. First, conditioned media (CM) from RRV RLI-infected SB28 cells was applied to CTLL-2 cells, a cytokine-dependent cytotoxic T cell line derived from C57BL/6 mice.^31^ The RLI secreted from infected tumor cells was able to simulate CTLL-2 growth to a level comparable to a recombinant IL15 control given at the same concentration (0.33 OD vs. 0.47 OD, p=0.01, **Fig. 2G**). Additionally, in a co-culture system, with SB28 tumor cells and naïve isolated mouse CD8^+^ T cells RRV RLI also significantly promoted the expression of the canonical T cell activation marker CD69 at a higher level than T cells cocultured with SB28 cells infected with RRV alone or uninfected SB28 tumor cells (p<0.005, **Fig. 2H**, gating strategy **fig. S1C**); confirming the functional ability of RLI to activate T cells. We also observed increased CD3^+^CD8^+^ T cell proliferation as indicated by Ki67 staining at 72 hours of incubation (p<0.05, **fig. S1D**) and improved CD8^+^ T cell persistence in culture (p<0.005, **fig. S1E).** While IL15 secreted by murine GBM cells transduced with RRV RLI stimulated the proliferation of cytotoxic T cells it did not increase the cytotoxic ability of T cells against tumor cells in co-culture when compared to RRV alone at multiple effector-to-target ratios (**Fig. 2H**). Subsequent co-culture assays with RRV RLI transduced SB28 cells and murine NK cells revealed the ability for RLI to significantly drive NK cell proliferation (p<0.005, **Fig. 2I**). RLI also promoted NK cell activation as evidenced by increased frequencies of double-positive CD69^+^GZMB^+^ NK cells (p<0.005, **Fig. 2J**) as well as CD69^+^ and GZMB^+^ single-positive NK cells (**fig. S1F and G,** gating strategy **fig. S1H**).

### RRV RLI treatment decreases intracranial tumor growth and prolongs survival in two poorly immunogenic murine models of glioblastoma

Having validated the stability and immunostimulatory effects of RRV RLI in culture, we next evaluated its therapeutic potential *in vivo* using intracranial SB28 and Tu2449 murine GBM lines implanted in syngeneic C57BL/6 and B6C3F1 mice (**Fig. 3A**). In the SB28 model, RRV RLI treatment significantly reduced tumor growth (**Fig. 3B**) and extended median survival compared to control (PBS or RRV alone, 19 vs. 55 days, p=0.0016), with long-term survival in 12% of treated mice (**Fig. 3C**). These findings were recapitulated and improved in the Tu2449 murine GBM model. In Tu2449 tumors, RRV RLI treatment led to pronounced tumor growth reduction on bioluminescent imaging (BLI) (**Fig. 3D**). Survival of RRV RLI treated Tu2449 mice was similarly significantly increased relative to control mice, with the majority of RRV RLI treated mice experiencing complete tumor regression and long-term survival (p=0.005, **Fig. 3E).** Rechallenge of mice cured from Tu2449 GBMs with contralateral intracranial injections of Tu2449 tumor cells demonstrated immunologic memory and rejection of injected cells with sustained survival (**Fig. 3F-G**). Blood samples from RRV RLI-treated SB28 mice showed no detectable RLI, confirming the localized effect of the treatment within the tumor immune microenvironment and the absence of systemic spread (**fig. S2A**).

**Fig 3.**
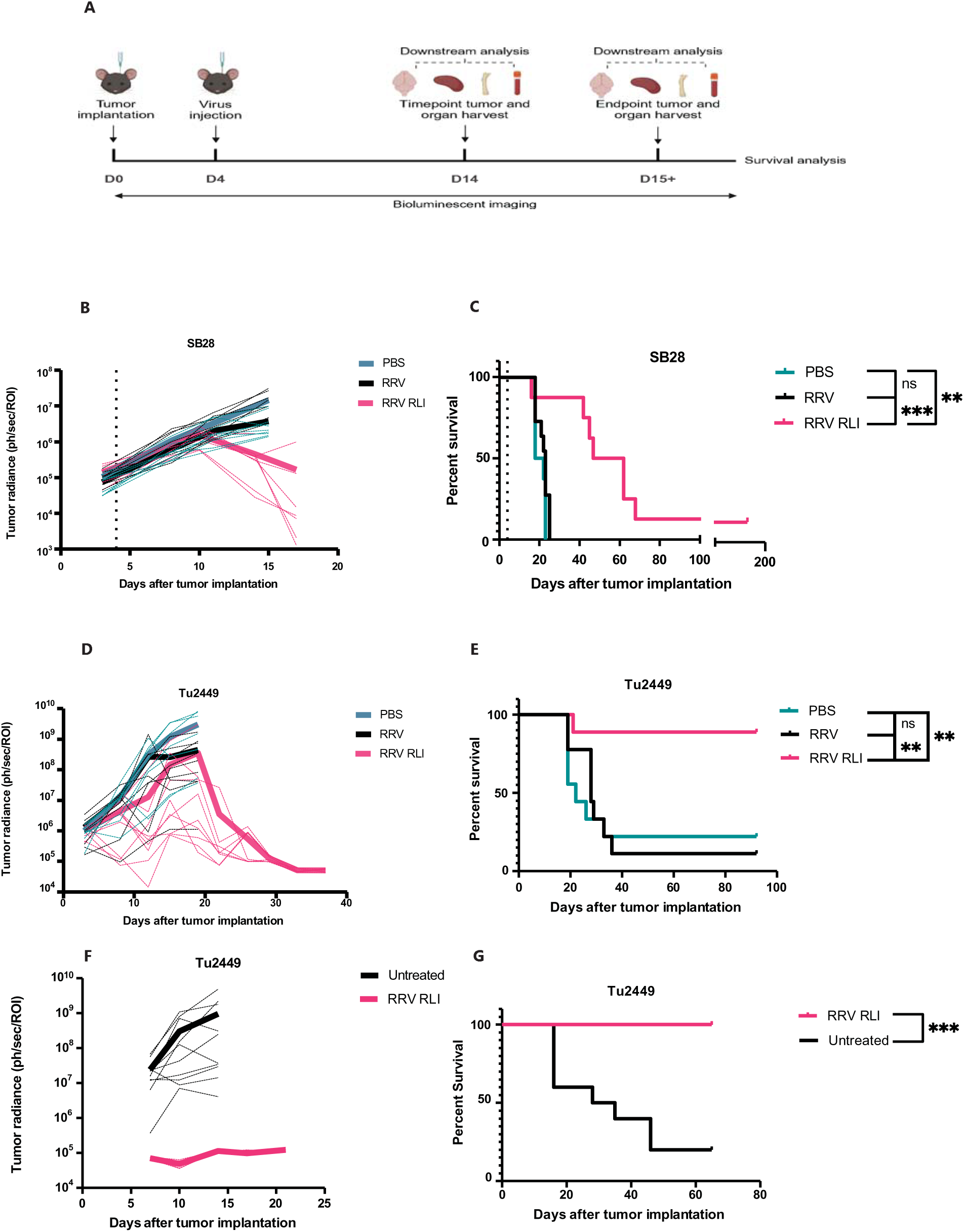
RRV RLI therapy suppresses tumor growth and prolongs survival in two poorly immunogenic murine glioblastoma models. (**A**) Schematic illustration of RRV RLI *in vivo* assessments. (**B**) RRV RLI treatment reduces average bioluminescent signaling 10 days post-tumor implantation in SB28 tumors. (**C**) RRV RLI significantly extends survival relative to PBS treated (median survival 54.5 days vs. 20 days, p<0.002, Log-Rank Mantel-Cox) and RRV treated (median survival 54.5 days vs. 23 days, p<0.001, Log-Rank Mantel-Cox) mice with SB28 tumors. (**D**) RRV RLI treatment decreases average bioluminescent signaling beginning 10 days after tumor implantation in Tu2449 tumors. (**E**) RRV RLI significantly prolongs survival relative to PBS treated (median survival undefined vs. 22 days, p<0.01, Log-Rank Mantel-Cox) and RRV treated (median survival undefined vs. 28 days, p<0.002, Log-Rank Mantel-Cox) mice with Tu2449 tumors. (**F**) No evidence of tumor growth is seen in bioluminescent imaging after contralateral intracerebral injection of 10,000 Tu2449 tumor cells in long-term previously tumor-bearing mice (**G**) Rechallenged mice display long-term survivorship relative to control (non-previously treated) mice (median survival undefined vs. 28 days, p<0.002, Log-Rank Mantel-Cox)

### RRV RLI induces significant anti-tumor modulation of the GBM immune microenvironment

To better understand the mechanisms behind RRV RLI’s therapeutic benefit, we transcriptomically analyzed the effect of RRV RLI treatment on immune cells in the SB28 microenvironment using the NanoString nCounter platform and a multiplex panel of 770 genes encoding markers of different immune cell types, common checkpoint inhibitors, and mediators of both the adaptive and innate immune responses. Differential gene expression analysis comparing RRV RLI-treated tumors to control tumors revealed that 14 days after tumor implantation and 10 days after RRV RLI treatment, tumors exhibited elevated transcription of the T cell modulating and co-stimulatory protein *Thy1* (fold change=5.2, padj=0.001); chemokine *Ccl8* (fold change=29, padj<0.001); cytotoxic T lymphocyte granule granzymes *Gzma* (fold change=81, padj<0.05) and *Gzmb* (fold change=51, padj<0.05); and T cell activation marker *Pdcd1* (fold change=42, padj<0.05), as well as reduced expression of *Cd276* (fold change=-4.3, padj<0.01), which encodes the immune checkpoint protein B7-H3 (associated with inhibiting cytotoxic T cell function and suppressing anti-tumor immune responses)^32,33^ (**Fig. 4A)**. Gene set enrichment analysis (GSEA) comparing RRV RLI to PBS treatment indicated that RRV RLI promoted the upregulation of T cell and NK cell functionality, antigen processing, and MHC pathways (**Fig. 3B**). Further interrogation of genes related to IL15 mediated functions, including T cell and NK cell functionality, antigen processing, and cytokine signaling revealed a diffuse upregulation in RRV RLI treated mice, including genes involved in the IL15 signaling pathway (**Fig. 4C-D**). Significantly upregulated genes of specific interest related to IL15 functions included those involved in MHC class I function and pathways (*H2-m3*, *H2-d1*, *Tap2*, *H2-t23*, *H2-k1*, *Psmb8*) and lymphocyte trafficking (*Ccr2, Ccr7, Ccr9, Cxr3*) (**Fig. 4C-D**). Gene network analysis uncovered a highly interconnected immune response with extensive interactions among genes involved in leukocyte cell-cell adhesion, mononuclear cell differentiation, and T cell differentiation, supporting the notion of a coordinated immune response elicited by RRV RLI treatment (**Fig. 4E**). Overall, RRV RLI-treated SB28 GBMs demonstrated a transcriptomic profile reflecting increased tumor-infiltrating T cells, NK cells, and dendritic cells relative to SB28 GBMs treated PBS (**Fig. 4F**). These findings align suggest that RRV RLI treatment promotes a robust and coordinated antitumor immune response.

**Fig 4.**
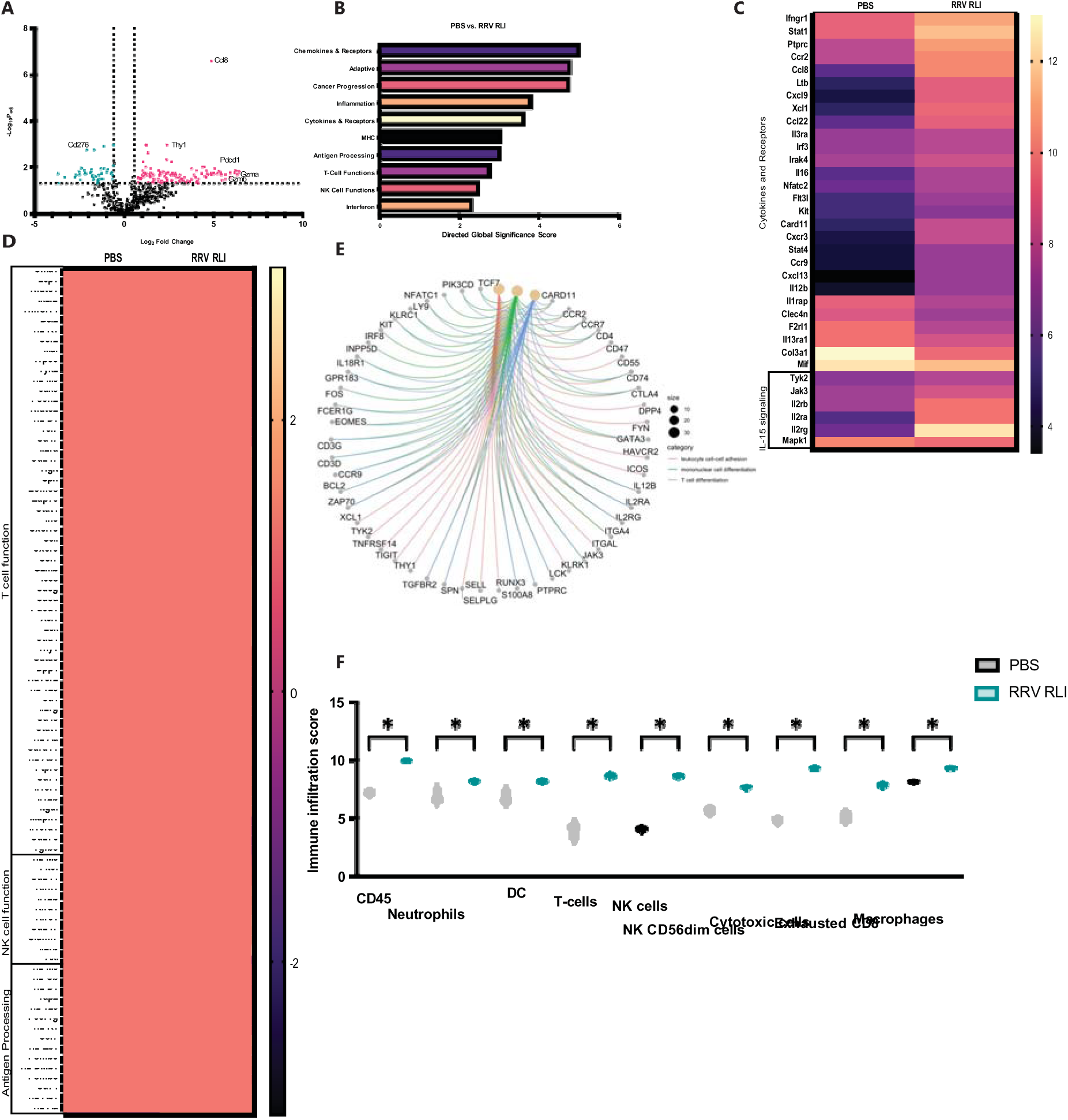
Comprehensive bulk transcriptomic analysis reveals extensive anti-tumor immune modulation induced by therapy. (**A**) Volcano plot showing differentially expressed genes between RRV RLI and PBS treatment groups at day 14 post-tumor implantation timepoint. (**B**) (**C**) Directed global significance scores of the top 10 upregulated gene pathways in RRV RLI treatment vs. PBS treatment. (**D**) Heat map detailing differential gene expression for genes related to T cell function, NK cell function, and antigen processing between RRV RLI and PBS treatment groups. (**E**) Gene network analysis of upregulated genes in RRV RLI treated glioblastomas compared to PBS-treated controls reveals a highly interconnected immune response network. (**F**) Violin plot demonstrating calculated immune infiltration scores between RRV RLI and PBS treatment groups at day 14 post-tumor implantation timepoint with significantly increased infiltration in RRV RLI across cell type (Welch’s t-test).

When comparing RRV RLI to RRV to isolate the effect of RLI, we similarly observed increases in pathways related to NK cell function, interferon, adhesion, T cell function, dendritic cell function, antigen processing, and MHC proteins (**fig. S2B**). Differential gene expression between RRV RLI and RRV-treated GBMs revealed increases in *Klra6* (modulator of NK cell function, promotes killing in the setting of downregulated MHC I, fold change=48, padj<0.001) and *Tnfrsf9* (4-1BB, powerful co-stimulatory molecule critical for enhancing T cell immune responses, fold change=160, padj<0.001)^34^ (**fig. S2C**). Upregulation of genes involved in mediating T and NK cell function or lymphocyte trafficking was also seen (e.g. *Ccr2, Cxcr3, Cxcl12*) (**fig. S2D**). When comparing RRV RLI-treated glioblastomas (GBMs) harvested at day 14 post-implantation versus at the endpoint, we observed a significant decline in most infiltrating immune cell populations, including T cells and NK cells. This suggests that the initial robust antitumor immune response induced by RRV RLI treatment lacks durability in mice that eventually failed to respond (**fig. S2E**). Additionally, mice treated with RRV alone showed a paucity of immune infiltration at the day 14 timepoint, implicating RLI as the primary driver of the therapeutic effect observed (**fig. S2E**).

### CD8^+^ and CD4^+^ T cells drive the antitumoral effects of RRV RLI in murine GBM

We then investigated the role of T cells and their subtypes in the antitumoral effects of RRV RLI in SB28 GBM treatment through antibody-mediated depletion studies (experimental layout, **Fig. 5A)**. As expected, with isotype control depletion alone, we continued to see a therapeutic benefit from RRV RLI treatment (p<0.005, **Fig. 5B-C**). However, when systemically depleting CD4^+^ and CD8^+^ T cells prior to tumor implantation and throughout the experiment after treatment, we observed complete abrogation of the RRV RLI therapeutic benefit (p=0.44, **Fig. 5D-E**). The efficacy of depletion was confirmed using the blood of tumor-bearing mice at 1 and 15 days after tumor implantation with reductions in CD3^+^, CD8^+^, and CD4^+^ populations to near undetectable levels (p<0.05 for all populations, **fig. S3A-D).** To better understand immune cell populations contributing to the RRV RLI therapeutic benefit we then investigated systemic CD8^+^ T cell depletion alone. While CD8^+^ T cell depletion resulted in 37.5% survivorship, there was no significant difference between RRV RLI and PBS control in tumor-bearing CD8^+^-depleted mice (p>0.1, **Fig. 5F-G**), implicating CD8^+^ cytotoxic T cells as being necessary for the therapeutic benefit seen with intratumoral RRV RLI treatment.

**Fig 5.**
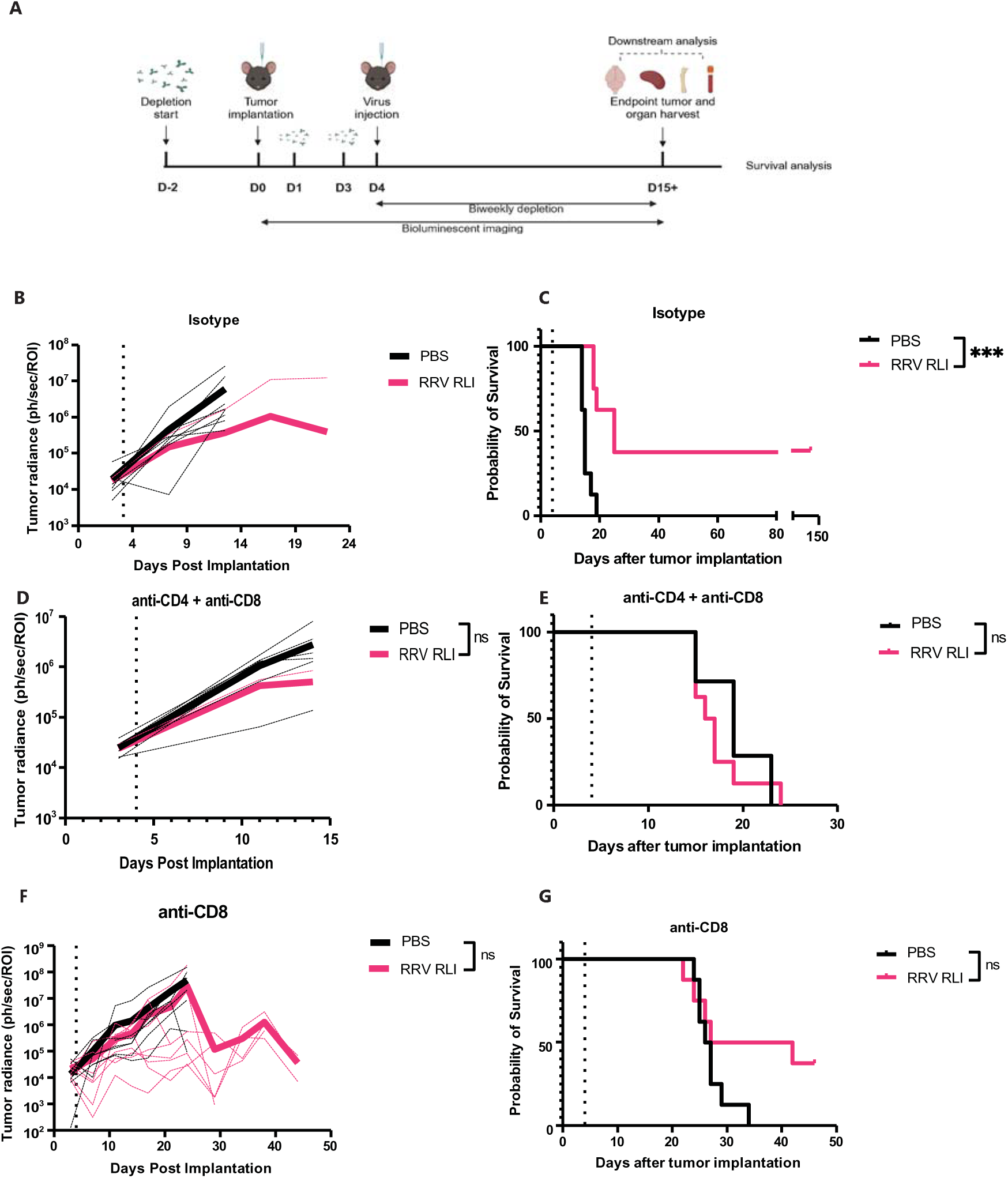
The survival benefit of RRV RLI therapy depends on both CD4**⁺** and CD8**⁺** T Cells. (**A**) Schematic illustration of RRV RLI *in vivo* assessment in T cell depletion experiments. (**B**) RRV RLI treatment reduces average bioluminescent signaling 9 days post-tumor implantation in SB28 tumors with isotype treatment. (**C**) In isotype antibody-treated mice bearing SB28 tumors, RRV RLI therapy significantly extended survival compared to PBS-treated controls (median survival 25 days vs. 15 days; p < 0.001, 37% long-term survivorship, Log-Rank Mantel-Cox test). (**D**) In anti-CD4 and anti-CD8 antibody-treated mice bearing SB28 tumors, RRV RLI therapy does not significantly change tumor growth on bioluminescent imaging (day 14, p=0.10, Welch’s t-test). (**E**) In anti-CD4 and anti-CD8 antibody-treated mice bearing SB28 tumors, RRV RLI does not significantly alter survival (median survival 16.5 days vs. 19 days, p=0.31, Log-Rank Mantel-Cox) when compared to PBS treated mice. (**F**) In anti-CD8 antibody-treated mice bearing SB28 tumors, RRV RLI therapy does not significantly change tumor growth on bioluminescent imaging (day 14, p=0.37, Welch’s t-test). (**G**) in anti-CD8 antibody-treated mice bearing SB28 tumors, RRV RLI does not significantly alter survival (median survival 34.5 days vs. 26.5 days, p=0.11, Log-Rank Mantel-Cox) when compared to PBS treated mice.

### Anti-PD1 therapy fails to overcome tumor escape from RRV RLI treatment

Given the dependence of RRV RLI on T cell-mediated antitumor immune responses, the observed upregulation of *Pdcd1* seen in our transcriptomic analysis, and the efficacy of anti-PD1 therapies in treating many types of cancer, we explored the therapeutic potential of combining RRV RLI with anti-PD1 blockade. Consistent with the known low immunogenicity of SB28,^35^ anti-PD1 treatment alongside PBS intracranial injection did not elicit a statistically significant improvement in bioluminescent imaging or survival in tumor bearing mice compared to isotype control (median survival 20 days vs. 23 days, p=0.132, **Fig. S3E-F**).

We then assessed whether inhibiting PD-1 signaling on circulating and intratumoral T cells via systemic anti-PD1 therapy could enhance therapeutic efficacy of intratumoral RRV RLI in the SB28 murine GBM model. While there was a trend towards decreased bioluminescent tumor signal (p=0.23, **fig. S3G**) and a higher percentage of long-term survivorship (20% vs. 10%, **fig. S3H**) this combination failed to yield a significant improvement in median survival or tumor regression when compared to RRV RLI with isotype control (median survival 43 days vs. 51.5 days, p=0.699, **fig. S3H**), suggesting that, while RRV RLI enhanced intratumoral immune cell infiltration and prolonged survival of GBM-bearing mice, PD-1 signaling alone was not a primary driver of treatment failure.

### Antitumoral efficacy of RRV RLI against murine GBM is potentiated by systemic chemotherapy

Given the importance of antigen quality and dominance in orchestrating an effective anti-tumor immune response,^36^ we then hypothesized that augmenting antigen presence in the TME by combining RRV RLI with systemically administered temozolomide (TMZ), a cytotoxic DNA-damaging alkylating chemotherapy that is standard of care treatment for newly diagnosed GBM, could potentiate the effects of T and NK cell infiltration induced by RRV RLI treatment (**Fig. 6A**). Indeed, the combination of systemic TMZ with intratumoral RRV RLI resulted in tumor remission in a significant proportion of treated mice with orthotopic SB28 GBM with improvement over RRV RLI monotherapy with vehicle control (median survival undefined vs. 48 days, p=0.0334) (**Fig. 6B**).

**Fig 6.**
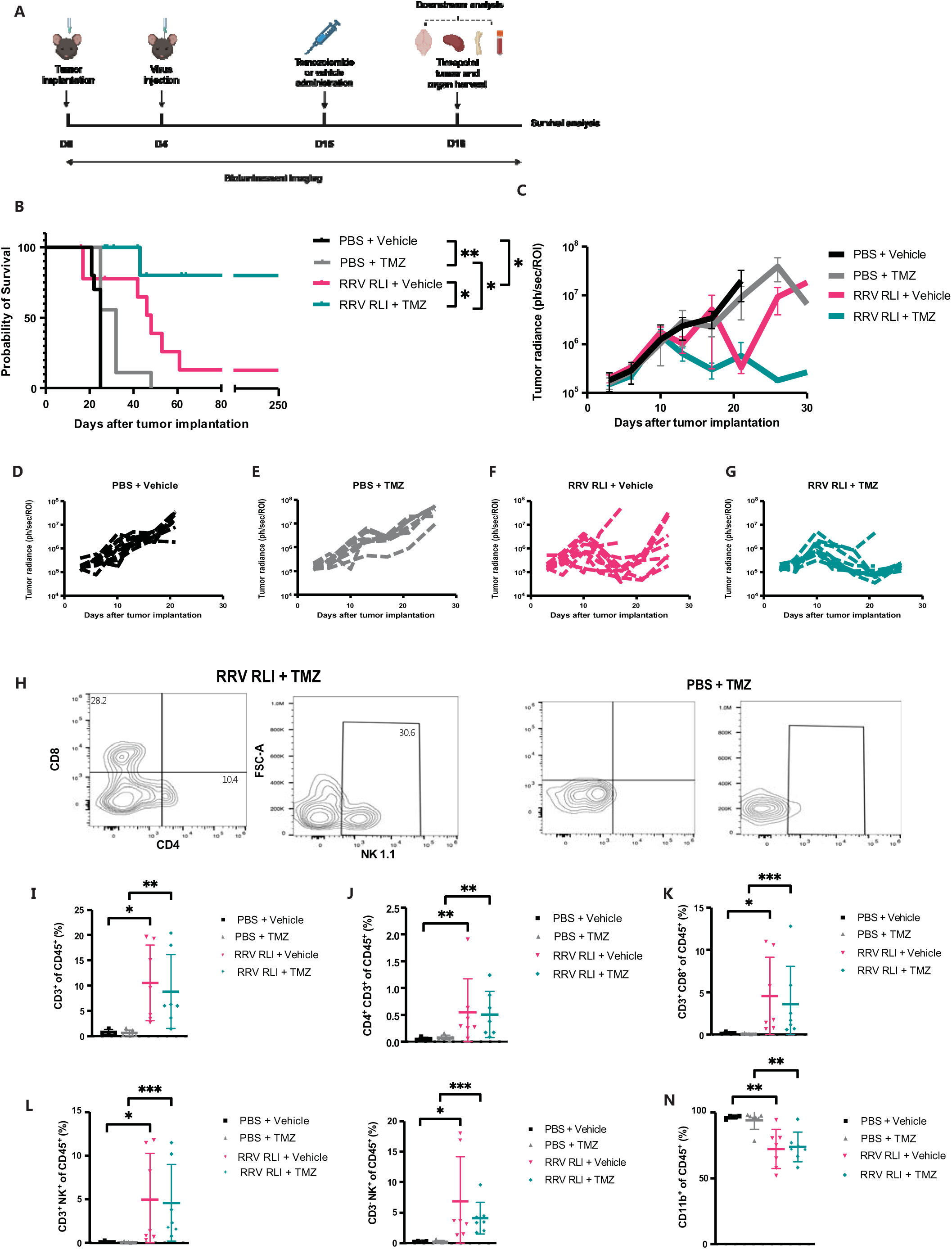
RRV RLI Synergizes with temozolomide chemotherapy to enhance survival. (**A**) Schematic illustration of RRV RLI *in vivo* assessment in temozolomide (TMZ) combination experiments. (**B**) RRV RLI + TMZ improves survival relative to RRV RLI + Vehicle (median survival undefined vs. 48 days; p<0.05, Log-Rank Mantel-Cox test) and PBS + TMZ (median survival undefined vs. 32 days; p<0.001, Log-Rank Mantel-Cox test). Mice reaching non-tumor endpoint were censored at that time. (**C**) Average bioluminescent tumor imaging. (**D-G**) Individual mouse bioluminescent imaging for all treatment groups. (**H**) Example flow cytometric gating for CD4 and CD8 T cells as well as NK cells at day 18 post-implantation timepoint. (**I**) Percent infiltrating CD3^+^ cells with significant differences between RRV RLI + TMZ and PBS + TMZ (p<0.01, Uncorrected Dunn’s test) as well as RRV RLI + Vehicle and PBS + Vehicle (p<0.02, Uncorrected Dunn’s test) (**J**) Percent infiltrating CD3^+^CD4^+^ cells with significant differences between RRV RLI + TMZ and PBS + TMZ (p<0.01, Uncorrected Dunn’s test) as well as RRV RLI + Vehicle and PBS + Vehicle (p<0.01, Uncorrected Dunn’s test) (**K**) Percent infiltrating CD3^+^CD8^+^ cells with significant differences between RRV RLI + TMZ and PBS + TMZ (p<0.01, Uncorrected Dunn’s test) as well as RRV RLI + Vehicle and PBS + Vehicle (p<0.05, Uncorrected Dunn’s test) (**L**) Percent infiltrating CD3^+^NK1.1^+^ cells with significant differences between RRV RLI + TMZ and PBS + TMZ (p<0.001, Uncorrected Dunn’s test) as well as RRV RLI + Vehicle and PBS + Vehicle (p<0.05, Uncorrected Dunn’s test) (**M**) Percent infiltrating CD3^−^NK1.1^+^ cells with significant differences between RRV RLI + TMZ and PBS + TMZ (p<0.001, Uncorrected Dunn’s test) as well as RRV RLI + Vehicle and PBS + Vehicle (p<0.05, Uncorrected Dunn’s test) (**N**) Percent infiltrating CD11b^+^ cells with significant differences between RRV RLI + TMZ and PBS + TMZ (p<0.01, Uncorrected Dunn’s test) as well as RRV RLI + Vehicle and PBS + Vehicle (p<0.01, Uncorrected Dunn’s test). All flow cytometry samples were collected at day 18 post-implantation timepoint.

This therapeutic synergy was further reflected in sustained reductions of bioluminescent signal in RRV RLI+TMZ combination cohort relative to RRV RLI + Vehicle and control groups (**Fig. 6C-G).** Flow cytometric analysis of intratumor immune cells (CD45^+^) revealed significantly enhanced infiltration of anti-tumoral T cell populations including CD3^+^, CD3^+^CD8^+^, and CD3^+^CD4^+^ cells in the RRV RLI treatment groups (RRV RLI + Vehicle or RRV RLI + TMZ) relative to controls (PBS+ Vehicle or PBS +TMZ) (**Fig. 6H-K**; additional CD3 gaiting strategy **fig. S4A).**

In line with the mechanism of RLI, further flow cytometric analysis demonstrated increased populations of NKT and NK cells in the RRV RLI treatment groups (**Fig. 6L and M**). Notably, there was no difference in T cell or NK cell populations between RRV RLI and RRV RLI TMZ treatment arms. The T cells and NK cells within the TME of RRV RLI treated mice exhibited high levels of Ki67 expression across CD3^+^, CD3^+^CD8^+^, and CD3^+^CD4^+^ cells with no difference between RRV RLI + Vehicle and RRV RLI + TMZ (**fig. S4B**).

We next focused on the behavior of myeloid populations in the TME. RRV RLI treated groups had significantly reduced infiltration of cells expressing the pan-myeloid marker CD11b with no difference between RRV RLI + Vehicle and RRV RLI + TMZ (**Fig. 6N**; gating strategy **fig. S4C**). Analysis of myeloid subpopulations revealed a decrease in macrophage (CD11b+ F4/80+) infiltration in RRV RLI+TMZ treated mice relative to mice receiving TMZ alone (p=0.01) though no significant differences were observed between RRV RLI treatment groups (**fig. S4D**). Conventional dendritic cell (CD11b^+^ CD11c^+^ MHCII^+^) infiltration did not significantly differ between treatment groups (**fig. S4E)**.

Flow cytometric analysis of peripheral blood from treated mice demonstrated a significant reduction in live CD45^+^ cells per 100 uL of blood in the TMZ-treated cohort compared to vehicle-treated animals, consistent with the lymphodepletion commonly observed in patients undergoing TMZ therapy. This reduction extended to CD3^+^ and CD3^+^CD8^+^ populations when comparing RRV-RLI + Vehicle and RRV RLI + TMZ (**fig. S4F-I**). These findings overall suggested that adding TMZ to intratumoral RRV RLI allowed the same levels of immune cells populations to infiltrate tumors despite the systemic myelosuppressive effects associated with TMZ treatment.

### Single cell sequencing of intratumoral leukocytes after treatment with RRV RLI and temozolomide reveals potent anti-tumor immune modulation

To understand how TMZ was potentiating the effects of RRV RLI without altering levels of the immune cell populations we interrogated by flow cytometry, we performed 5’ single-cell sequencing of CD45^+^ leukocytes, isolated by FACS (fluorescence-activated cell sorting) from SB28 GBMs treated with intratumoral RRV RLI and/or systemic temozolomide. After processing, a total of 28,125 cells were annotated into 17 different clusters (**Fig. 7A**). T and NK cells were then relabeled into 12 phenotypic clusters based on predetermined markers (**Fig. 7B**), with the distribution of critical gene markers provided in **Fig. S5A**. RRV RLI treatment groups exhibited higher levels of T and NK cell infiltration as compared to PBS control groups, with marked expansions in specific clusters including NKT cells, NK cells, and CD8^+^ T cell populations (**Fig. 7C**).

**Fig 7.**
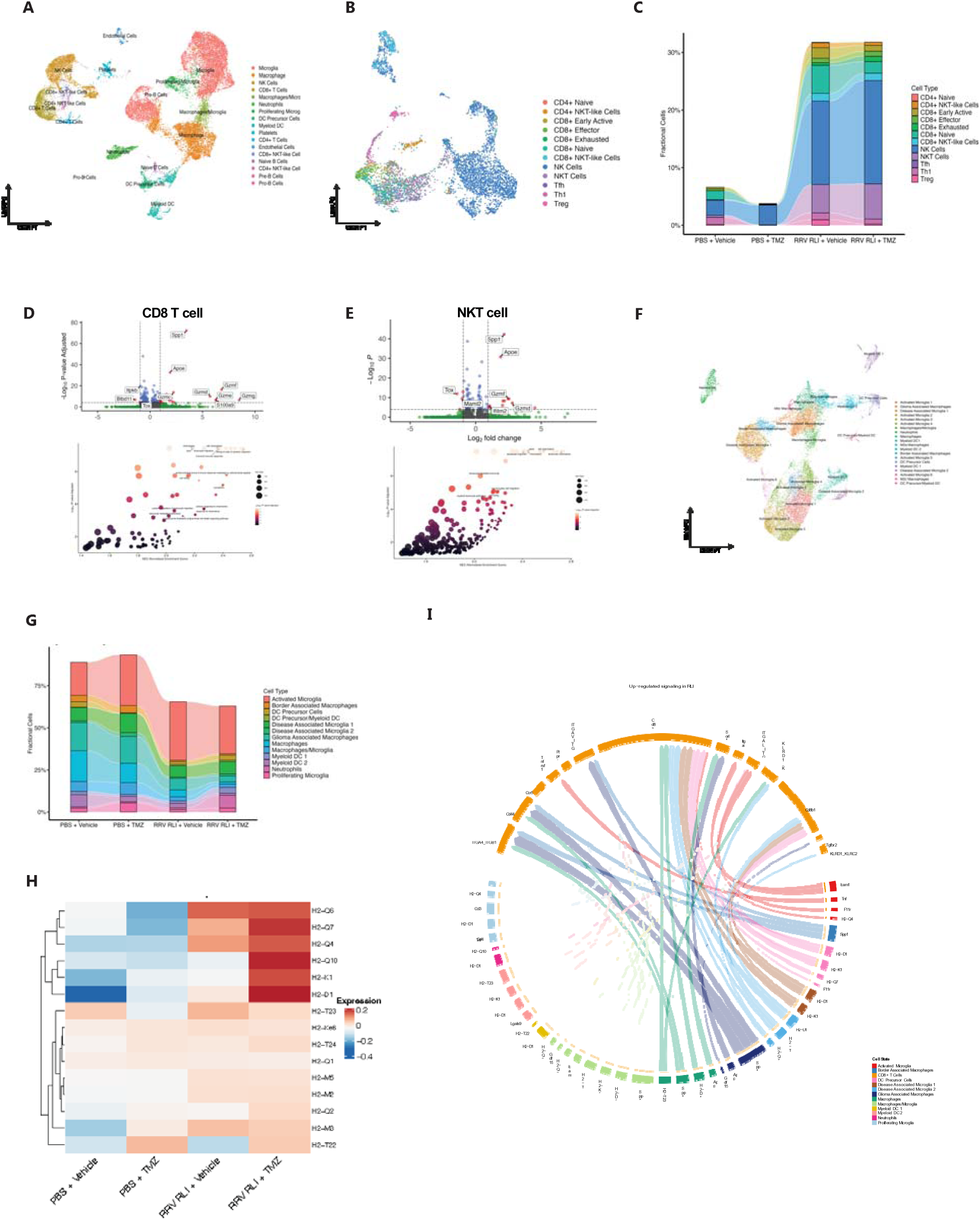
Single cell RNA sequencing reveals cellular mechanisms driving therapeutic efficacy and its enhancement with TMZ. (**A**) Single-cell RNA sequencing UMAP of all sequenced transcriptomes from CD45^+^ cells isolated from SB28 tumor-bearing mice, demonstrating myeloid, T, and NK cell clusters. (**B**) T and NK cell focused UMAP including all samples. (**C**) Alluvial plot demonstrating changes in gross T and NK cell infiltration between samples. (**D**) Volcano plot demonstrating CD8 T cell differential gene expression in RRV RLI + TMZ vs. RRV RLI + Vehicle with additional GSEA. (**E**) Volcano plot demonstrating NKT cell differential gene expression in RRV RLI + TMZ vs. RRV RLI + Vehicle with additional GSEA. (**F**) Myeloid cell-focused UMAP including all samples. (**G**) Alluvial plot demonstrating changes in gross myeloid cell infiltration between samples. (**H**) Heat map of MHC class I expression in myeloid populations across samples highlighting increased expression in RRV RLI + Vehicle and enhancement in RRV RLI + TMZ. (**I**) Cell chat analysis of incoming signaling in RRV RLI + Vehicle vs. PBS + Vehicle.

Differential gene expression analysis of tumor-infiltrating CD8^+^ T cells between RRV RLI + TMZ and RRV RLI + Vehicle demonstrated upregulation of genes indicative of cytotoxic immune activity and antitumor immunity including *Gzmb* (log2FC=0.67, padj=2.13e-12) *, Gzmc* (log2FC=2.23, padj =1.5e-14)*, Gzmd* (log2FC=5.72, padj =1.32e-10)*, Gzme*(log2FC=6.27, padj=2.53e-7)*, Gzmf* (log2FC=6.81, padj=4.72e-17), and *Gzmg*(log2FC=8.50, padj=4.14e-7) (**Fig. 7D**). Additionally, there was notable downregulation of *Tox* (log2FC=-1.0, padj=4.45e-5), indicative of reduced T cell exhaustion and decreased expression of inhibitory receptors (**Fig. 7D**).^37^ Further gene set enrichment analysis (GSEA) revealed enrichment of biological processes related to chemotaxis, cell killing, and response to chemokines in RRV RLI + TMZ as compared to RRV RLI + Vehicle (**Fig. 7D**). Similar trends in gene expression and GSEA were seen when comparing NKT cells between these groups (**Fig. 7E**). ^38^

Attention was then turned to myeloid populations across the different treatment groups, identifying 20 different myeloid sub-clusters using scType^38^ and expression of canonical marker genes (**Fig. 7F**, **fig. S5B**).^39–41^ As observed in flow cytometric analyses, myeloid populations were a lower fraction of total CD45^+^ cells in RRV RLI treatment groups, with a disproportionate decrease in the macrophage population when compared to PBS groups (**Fig. 7G**). However, MHC class I gene expression was increased in RRV RLI treatment groups, with even higher expression overall in the RLI + TMZ combination. This increase was particularly evident in genes including *H2-q6, H2-q7, H2-q4, H2-q10, H2-k1, H2-d1* (**Fig. 7H**). Specific myeloid cell populations with increased MHC class I gene expression varied but included myeloid DC 1 and 2 cells, neutrophils, and precursor myeloid DC cells (**fig. S5C-D**). These findings supported the hypothesis that TMZ-induced cell death and antigen release led to enhanced antigen-presentation in the GBM TME.

To assess potential interactions between these gene expression changes we conducted a ligand-receptor analysis using CellChat. Comparing CD8^+^ T cells between RRV RLI + Vehicle and PBS + Vehicle identified increased receptor-ligand signaling dominated by antigen presentation signaling primarily through MHC class I proteins from multiple myeloid cell populations (including neutrophils, myeloid DC 1, myeloid DC2, and macrophages/microglia) to the CD8^+^ T cell population (**Fig. 7I**) suggestive of cross-presentation of antigens. This antigen presentation signaling was further amplified when comparing CD8^+^ T cells between RRV RLI + TMZ vs. RRV RLI + Vehicle (**fig. S5E**). Similarly, enhanced signaling pathways related to migration and immune cell infiltration (e.g., Spp1 signaling) were observed in both comparisons (**Fig. 7I and fig. S5E**).

### T cell receptor sequencing suggests tumor antigen-specific T cell responses induced by RRV RLI treatment

To further elucidate the impact of each treatment on the T cell receptor (TCR) repertoire on tumor-infiltrating T cells, TCR sequencing was performed on tumor-infiltrating T cells. Comparison of TCR repertoire between treatment groups revealed over double the percent recurrent TCRs sequenced in the RRV RLI + TMZ treatment group vs. the RRV RLI + Vehicle group (15.3% vs. 7.2% **fig. S6A**), suggestive of clonal expansion and an antigen-driven immune response.^42^ This was further supported by a higher percentage of the TCR repertoire space being occupied by recurrent clonotypes in tumors treated with RRV RLI + TMZ vs. RRV RLI alone (**fig. S6B**), indicating that combining RRV RLI and TMZ led to an antigen-specific immune response.

T cell receptor (TCR) clustering was performed using CDR3 beta-chain sequences from all samples. We found that 1,810 out of 1,813 TCRs were within 80 distance units of each other (**fig. S6C**). The top 10 clusters accounted for 91.7% of the sequenced TCRs (1,659 out of 1,813 TCRs), and each of these clusters included TCRs from a PBS-treated group (non-virus exposed, **fig. S6D and E**).

We then assessed the similarity between our TCR sequences and those specific to pathogens and murine leukemia virus (MLV) as reported in the McPAS-TCR database.^43^ In total, 42 out of 1,813 beta chain TCRs (2.3%) were within 18 distance units of a pathogen-specific TCRs from McPAS-TCR. Notably, none of the TCRs reactive to the envelope protein of MLV were within 100 distance units for paired chains or 18 distance units for beta chains in our samples.

Subsequently, we performed exact matching on the beta chain TCR data. There were 45 exact matches across all TCR sequences. Importantly, no exact matches were found in our data for beta chains of TCRs reactive to MLV envelope proteins. We identified a total of 24 exact matches corresponding to pathogen-specific TCRs, which mapped to herpes simplex virus type 1 (HSV-1), influenza virus, and murine cytomegalovirus (mCMV). Together, these findings suggested that the TCRs we had identified were part of a tumor-specific response enhanced by RRV RLI and RRV RLI+TMZ rather than an antiviral response triggered by RRV RLI.

## Methods

### Study design

The objective of this study was to assess the therapeutic potential of RRV RLI and to elucidate the underlying biological mechanisms driving its efficacy. For in vivo experiments, 8-12 week-old mice were utilized, with 6 to 10 mice per group, ensuring adequate statistical power based on preliminary data. Mice were randomized into treatment arms following tumor implantation, with randomization based on bioluminescent imaging to equalize the average starting tumor sizes across groups. All experiments adhered to Institutional Animal Care and Use Committee (IACUC) guidelines, with predetermined survival endpoints applied. Mice reaching non-tumor endpoints were censored from survival analyses.

### Construction of RRV RLI

Gibson assembly cloning was used to place a codon- and stability-optimized genetic sequence for RLI^44^ into pAC3-P2A-yCD. A T2A cleavage peptide sequence was positioned at the C-terminus of the viral envelope protein followed by the RLI sequence (**Fig. 2A**).

### Cell lines and culture

Human embryonic kidney 293T (Lenti-X cells, purchased from Takara, Inc.), murine glioblastoma Tu2449 (generously provided by Dr. Noriyuki Kasahara, University of California, San Francisco), and human glioblastoma U87 (generously provided by Dr. Noriyuki Kasahara, University of California, San Francisco) were cultured in Dulbecco’s modified Eagle’s medium (DMEM) supplemented with 10% fetal bovine serum (FBS) and 1X Gibco GlutaMAX (Gibco, Inc.). Murine glioblastoma SB28 (generously provided by Dr. Hideho Okada, University of California, San Francisco) were cultured in RPMI 1640 medium supplemented with 10% FBS, 2% Gibco GlutaMAX, 1% non-essential amino acids (Gibco, Inc.), 1% HEPES (Gibco, Inc.), 1% Penicillin-Streptomycin (Gibco, Inc.), and 0.1% beta-mercaptoethanol. Cells were screened bimonthly for mycoplasma and validated every six months by Short Tandem Repeat (STR) analysis at the University of California Cell Culture Facility.

### Viral production and concentration

Virus was produced via transient transfection and the use of stable producer cell lines. For transient transfection protocols, reverse transfection utilizing Fugene HD (Promega) and 10 micrograms of viral plasmid DNA was implemented. Stable U87 producer cell lines were also utilized for viral production. Virus containing supernatant was collected at approximately 36-48 hours after initial transfection. For *in vivo* studies virus was concentrated using column-based retrovirus purification with buffer exchange to phosphate buffer solution (PBS) (Bioland Scientific LLC.). Functional viral titers in transducing units per mL (TU/mL) were determined via staining for the viral gag-protein and flow-cytometry.

### In vitro viral replication and stability

Viral replication and stability were performed in a similar manner to previous studies.^45^ In brief, specific multiplicities of infection (MOI) were added to tumor cells in vitro and allowed to replicate over time, with transduction levels measured at regular intervals via staining for the gag viral protein and flow cytometry. Azidothymidine, which inhibits viral spread, was utilized for control groups. For viral stability, an MOI of 0.01 was added to tumor cells and allowed to spread to 100% transduced cells at 14 days. The resulting supernatant was then placed on tumor cells at an MOI of 0.01 and the protocol was repeated for a total of 8 weeks. Genomic DNA was then isolated using the Monarch® Genomic DNA Purification Kit (NEB Biolabs). Polymerase chain reaction with primers crossing the RLI insert were used to determine stability of the construct. Primer sequences were as follows: FWD: ggaccttgcattctcaatcgattgg REV: cccctttttctggagactaaataa (**fig. S1B**).

### RLI production and function

RLI production levels were determined using supernatant from 100% infected SB28 tumor cells. RLI levels in the supernatant was measured via Human IL15 Quantikine enzyme-linked immunosorbent assay (R&D Systems). RLI function was determined using the CTLL-2 proliferation assay similarly to previous studies.^46^ In brief, RLI from transduced cells was added to cytokine-starved CTLL-2 cells with proliferation of these cells subsequently measured via Colorimetric MTS Assay (Promega). Co-culture assays were conducted by incubating SB28 tumor cells with freshly isolated NK cells (BioLegend MojoSort™ NK Cell Isolation Kit) or CD8+ T cells (BioLegend MojoSort™ CD8 T Cell Isolation Kit) from C57BL/6 spleens. Tumor and effector cells were co-cultured at varying tumor-to-effector cell ratios and conditions. Activation markers were measured via flow cytometric analysis. Cytotoxic activity was measured using the xCelligence real-time cell analysis system.

### Animal studies

Animal experiments were approved by UCSF IACUC (approval #AN105170-02). C57BL/6 and B6C3F1 mice (8–12 weeks old) were obtained from Jackson Laboratories and housed at the University of California, San Francisco. Experiments utilizing Tu2449 and SB28 murine glioblastoma cell lines were conducted under comparable conditions. All cell lines expressed luciferase to facilitate bioluminescent imaging. On day 0, 10,000 tumor cells were implanted intracranially using a stereotactic frame at the following coordinates relative to the bregma: anteroposterior (AP), 0 mm; mediolateral (ML), 1.2 mm; and dorsoventral (DV), 3.5 mm. Mice were imaged on day 3 or day 4 post-tumor injection to establish pretreatment bioluminescent baselines, after which they were randomized into treatment groups. On day 4 post-tumor implantation, mice were injected with 2.5 × 10^5 TU of either RRV RLI or control RRV, depending on the assigned treatment group.

For the anti-PD1 combination experiments, mice were administered either anti-PD1 antibody (RMP1-14, BioXcell) or isotype control (BioXcell) at a dose of 200 µg, delivered via intraperitoneal injection on day 7 and day 24 post-tumor implantation, with subsequent dosing every other day for a total of four doses. In the TMZ combination experiments, mice were treated with temozolomide (TMZ) at a total dose of 400 mg/kg, administered over three days (diluted in 10% DMSO, T2577 Sigma Aldrich) starting on day 15 post-virus injection, or with vehicle control (10% DMSO).

For T cell depletion studies, depletion antibodies were administered two days prior to tumor implantation and subsequently continued biweekly post-implantation. A total of 400 µg of anti-CD8 antibody (200 µg YTS 169.4, BioXcell; 200 µg 53-6.7, BioXcell) and 200 µg of anti-CD4 antibody (GK1.5, BioXcell) were administered. Control mice received 600 µg of isotype antibody (LTF-2, BioXcell). Depletion was confirmed on days 1 and 15 after tumor implantation. Survival data were plotted using the Kaplan-Meier method, with statistical comparisons between groups performed using the Log-rank test (GraphPad Prism 9).

### Analysis of immune alterations

Mice were sacrificed at endpoint, day 14 post-tumor injection timepoint (non-TMZ studies), or day 18 timepoints (TMZ studies). The following tissues were harvested: spleen, bone marrow, blood, and brain tumor. Brain tumors were minced and digested in collagenase type IV (Thermo Fisher Scientific, #17104019) and Deoxyribonuclease I (Worthington Biochemical Corporation) solutions while agitated at 37°C. Tumor suspensions were subsequently filtered through 70 µm filters, and red blood cells were lysed using Ammonium-Chloride-Potassium (ACK) lysing buffer (Lonza). Spleens were dissociated through 40 µm filters and similarly subjected to ACK lysis, as was the bone marrow.

Flow cytometric analysis was then performed on processed tissues. A detailed list of antibodies can be found in Supplemental Table 1. Briefly, cells were first blocked with mouse Fc block in PBS containing 2% bovine serum albumin (BSA). After Fc blocking, cells were washed and stained with Zombie Aqua fixable viability dye (BioLegend, #423101) in PBS. Following viability staining, cells were washed again and stained for surface markers in PBS with 2% BSA. Intracellular marker staining was performed using the eBioscience™ Foxp3/Transcription Factor Staining Buffer Set (Thermo Fisher Scientific, #00-5523-00). Data acquisition was conducted using an Attune NxT Flow Cytometer (Thermo Fisher Scientific), and flow cytometry data were analyzed using FlowJo software.

### Nanostring multiplex transcriptomic analysis

RNA was extracted from murine tumor single-cell suspensions using the RNeasy Mini kit (Qiagen) and stored at −80°C until further use. RNA quality and quantity were assessed using a bioanalyzer. For each sample, 100 ng of RNA was hybridized with the Nanostring Mouse PanCancer Immune Profiling codeset for 18 hours. A 30 µL aliquot of the reaction was loaded into the nCounter cartridge and processed on the nCounter SPRINT Profiler. Quality control and raw data alignment were conducted using nSolver (Nanostring).

Differential gene expression analysis was performed using the DESeq2 package in R, followed by pathway expression analysis as defined by the KEGG 2019 Human database, utilizing the Enrichr pipeline. Simultaneously, raw files were analyzed on the Rosalind online platform (OnRamp Bio) to calculate normalized gene expression counts, determine significance of differentially expressed genes, and derive cell type scores. Visualizations, including heatmaps and volcano plots, were generated using GraphPad Prism 10.

### Mouse scRNA sequencing and immune cell profiling analysis

Brain tumors were processed as previously described on day 18 after tumor implantation. CD45^+^ cells were isolated via fluorescence activated cell sorting. 5’ scRNA-seq with TCR sequencing was carried out using the 10X Genomics platform and per manufacturer instructions. Sequencing was performed on the Illumina Novaseq 6000.

Single-cell FASTQ files, along with paired TCR V(D)J immune profiling FASTQ files, were aligned using CellRanger (version 7.2, 10X Genomics). The resulting expression matrices were imported into RStudio (version 4.3.2) for downstream analysis. Standard pre-processing was conducted using Seurat (version 5.0.3). The dataset included 2,928 cells in the PBS condition, 8,716 cells in the TMZ condition, 6,225 cells in the RRV RLI + Vehicle condition, and 10,872 cells in the RRV RLI + TMZ condition. Cells expressing more than 200 genes with less than 10% mitochondrial gene expression were retained for further analysis.

Samples were integrated using the standard Seurat v5 workflow. In total, 28,125 cells were assigned across 25 clusters. Cell types were identified using a combination of suggested labels from scType and marker gene expression based on literature review. Alluvial bar graphs showing sample composition by cell type were created using ggalluvial (version 0.12.5).

The integrated samples were divided into lymphocyte and myeloid subsets based on cluster identity. The lymphocyte subset included cells labeled as NK Cells, CD4+ T Cells, CD8+ T Cells, CD8+ NKT-like Cells, and CD4+ NKT-like Cells, while the myeloid subset contained Macrophages, Microglia, Proliferating Microglia, Macrophages/Microglia, Myeloid DCs, Neutrophils, and DC Precursor Cells. Cells in the lymphocyte subset were relabeled according to phenotypes determined by ProjecTILs (version 3.3.0). Cells assigned an “NA” label were presumed to be NK cells, and cells expressing both Cd3e and Klrb1c above the 20th percentile were classified as NKT cells.

Volcano plots were generated using EnhancedVolcano (version 1.2.0). Gene Set Enrichment Analysis (GSEA) was performed using ClusterProfiler (version 4.10.1). Ligand-receptor interaction analysis was conducted with CellChat (version 2.1.2).

T cell V(D)J sequencing data was analyzed using immunarch (version 1.0.0), and unique clonotypes were identified based on paired TRA and TRB gene sequences. By sample, 66 T cells were sequenced in the PBS condition, 15 in the TMZ condition, 755 in the RLI condition, and 1,052 in the RLI + TMZ condition. Immunarch was also used to find matching TCR CDR3 beta sequences in the McPAS databass. TCR distance analysis was performed using the tcrdist3 package (v0.2.2) in Jupyter notebook (python 3.12.7). All single cell sequencing data analyzed in this study can be accessed from the Gene Expression Omnibus repository, accession code GSE278988.

### Human bulk RNA-sequencing analysis

To evaluate the gene expression of IL15 across different cancer types, the TCGA PanCancer Atlas from Hoadley et al. was accessed via cBioPortal (cbioportal.org) for IL15 RSEM scaled estimates in available solid tumor samples.^47^ To investigate differences according to magnitude of IL15 expression in glioblastoma, the TCGA cohort of primary glioblastoma samples was stratified into high and low IL15 expression at the 80th percentile after extracting HTSeq counts via TCGAbiolinks and converting into counts per million (CPM) via edgeR.^48^ Differentially expressed genes were evaluated between high and low IL15 glioblastoma patients via DESeq2 and defined as genes with a Benjamini–Hochberg adjusted p-value < 0.05 and a |log2Fold Change| > 2. Gene ontology enrichment analysis was performed on overexpressed genes in high IL15 glioblastoma patients to identify the overrepresented biological processes, cellular components, and molecular functions in those tumors. To quantify and compare the immune cell infiltration between high and low IL15 glioblastoma patients, RSEM scaled estimates were converted into transcripts per million (TPM) and processed through the CIBERSORT web application (https://cibersortx.stanford.edu/), a deconvolution algorithm that infers the proportion of 22 types of tumor-infiltrating immune cells from bulk RNA-sequencing data.^49^ Degree of immune cell infiltration was compared using the Mann-Whitney U test. A two-tailed p-value < 0.05 was used as the threshold for statistical significance.

### Human scRNA and immune cell profiling analysis

Differences in transcriptomic expression were investigated by analyzing single cell RNA sequencing data from four previously published GBM resection specimens.^29^ Clusters and cell identities largely mirrored those identified in the original study; however, immune cells were further subdivided to refine cell-type classification. This subclassification of immune cells was achieved using labels suggested by SingleR and marker gene expression identified through literature review. Samples were classified as IL15 High or IL15 Low based on standard analysis of IL15 gene expression levels. Differential gene expression was visualized using volcano plots generated with EnhancedVolcano. Gene ontology analysis was conducted using ToppGene, and cellular communication via ligand-receptor interactions was assessed using CellChat.

### Statistical analysis

Statistical analyses were performed using Prism 10 (GraphPad). Specific study parameters and statistical methods are detailed within the figure legends. Comparisons between two groups were conducted using a two-tailed Student’s t-test. For comparisons among multiple groups, a one-way ANOVA was employed, followed by Fisher’s LSD post-hoc test for pairwise comparisons, assuming a Gaussian distribution and equal standard deviations. For non-parametric comparisons, the Kruskal-Wallis test with Dunn’s post-hoc test was applied. Kaplan-Meier analysis was used for in vivo survival studies, with differences assessed via the log-rank (Mantel-Cox) test. Outliers were removed based on Grubbs’ test.

## Discussion

The IL15 superagonist RLI is a potent immunostimulatory agent with the ability to enhance an anti-tumor immune response through T and NK cell modulation. Our study evaluates RRV RLI as a viral immunotherapy for the treatment of GBM. We demonstrate that tumor cells infected with RRV RLI produce functional RLI with canonical functions such as supporting T and NK cell proliferation. Single agent intratumoral RRV RLI treatment resulted in significantly improved survival in the immunosuppressive SB28 model and long-term survival with immunologic memory to contralateral orthotopic rechallenge in the Tu2449 GBM model. RRV RLI also synergizes with GBM standard of care in the form of TMZ chemotherapy to provide a long-term survival benefit in the SB28 model. RRV is a highly translatable vector platform with a strong clinical safety profile in patients even when delivered intravenously.^30^ This work establishes RRV RLI as a novel and translatable viral cancer immunotherapy underscoring its potential for clinical application in humans.

IL15 is regarded as a promising immunocytokine for cancer immunotherapy.^50^ While other cytokines, such as IL-2, promote T cell growth, they also act as a ‘double-edged sword’ by concurrently upregulating T-regulatory cells and triggering activation-induced cell death (AICD) or capillary leak syndrome. In comparison, IL15 supports the proliferation and activation of CD8 T cells (including memory phenotype), NKT cells, and NK cells.^51–54^ Moreover, IL15 induces pro-inflammatory changes within the tumor microenvironment and has demonstrated synergy when combined with other immune and chemotherapeutic agents. ^55,56^

Local delivery of immunotherapies directly to the tumor microenvironment mitigates the dose-limiting toxicities often associated with systemic administration. In the brain, this approach also circumvents the blood-brain barrier (or blood-tumor barrier), a major obstacle for many systemically administered treatments. In this study, we aimed to transform glioblastoma tumor cells into biofactories for RLI through delivery via RRV, a virus that selectively spreads in dividing cells without inherently killing host cells. Our rationale was that this strategy would create a portion of tumor cells secreting RRV RLI and a field effect across the tumor with the goal of *in situ* tumor vaccination.

We observed that local delivery and tumor-mediated secretion of RLI in intracranial tumors produced outcomes in line with or superior to those observed with systemic administration of RLI in other cancer types or ALT-803, an alternative IL15 superagonist, in GBM.^57–59^ An advantage of our study is that our findings were in two syngeneic GBM models with best-in class translatability with regards to the investigation of immunotherapies.^35^ Directed tumor delivery of RRV RLI led to a significant survival benefit in both the SB28 and Tu2449 GBM models. Notably, treated Tu2449 mice exhibited over 90% long-term survivorship and developed immunologic memory upon intracranial rechallenge, effectively demonstrating successful vaccination against tumor cells. We attribute the differential response to single-agent RRV RLI treatment between the two models to variations in their underlying tumor immunogenicity. While Tu2449 is also poorly immunogenic relative to most murine GBM models, it does exhibit higher T cell infiltration and a higher (but still low) baseline rejection rate compared to the SB28 model, indicating increased immunogenicity. SB28 being one of the least immunogenic models makes it more representative of human GBM, characterized by low T cell infiltration and a low tumor mutational burden.^35,60^ Impressively, RRV RLI treatment alone significantly increased CD8 T cell, NKT cell, and NK cell infiltration in treated SB28 mice from near undetectable levels. We believe these findings are promising for the treatment of human GBM, where CD8+ T cell infiltration is estimated to be under 5% of tumor-infiltrating immune cells, similar to the SB28 model.^61^ RRV RLI treatment also induced broad transcriptomic inflammatory changes within the tumor microenvironment and upregulated antigen-presenting pathways. These findings align with existing literature highlighting the role of IL15 in the maturation and function of antigen-presenting cells.^62^ Collectively, this manipulation of the local tumor environment enhanced the immunogenicity of an otherwise immunologically silent tumor.

Given the increased infiltration of both T cell and NK cell populations in the tumor microenvironment, we sought to examine the critical mediators of the RRV RLI treatment response. Simultaneous depletion of both CD4^+^ and CD8^+^ T cells as well as CD8^+^ T cells alone abrogated the survival benefit imbued by RRV RLI treatment, indicating a reliance on those T cell populations for treatment efficacy, despite increased NK cell infiltration in treated tumors. This is in line with previous work using ALT-803 in GBM which saw a reduction in treatment efficacy with CD4^+^ or CD8^+^ T cell depletion, but not with NK cell depletion.^63^ The role of NK cells in mediating anti-tumor efficacy in response to systemic IL15 immunotherapy has varied, with some cancer models demonstrating a dependence on NK cells^64^ and others with a reliance on tissue-resident CD8^+^ T cells. ^59,65^ Interestingly, CD8 depletion did not significantly affect overall survival but did result in a subset of long-term survivors among RRV RLI-treated mice, indicating a heterogeneous response to RRV RLI in this context, with the lack of long-term survivors in mice with simultaneous CD4 and CD8 depletion potentially suggesting a dynamic compensatory role for CD4^+^ T cells and NK cells in the RRV RLI response in the absence of CD8^+^ T cells.

An inherent challenge to the translation of immunotherapies to GBM is understanding interactions with standard of care, especially temozolomide which causes myelosuppression as a known dose-limiting toxicity.^66,67^ Interestingly, the combination of RRV RLI and temozolomide worked synergistically to provide a significant survival benefit compared to either treatment alone. Flow cytometric analysis of the tumor microenvironment revealed no gross differences in immune cell infiltration between the RRV RLI monotherapy and the combination therapy with temozolomide, highlighting the sustained presence of tumor-infiltrating T cells despite systemic myelosuppression as indicated on flow cytometry of peripheral blood samples. This has important implications for clinical translation given the widespread use of TMZ in patients. 5’ single cell RNA sequencing of the treatment groups also revealed similar immune cell infiltration, in congruence with the flow cytometry-with no difference between RRV RLI alone and with TMZ. Interestingly, RRV RLI treatment elicited enhanced activation of T cells and NKT cells, evidenced by the upregulation of cytotoxicity-associated genes such as Gzmb and the downregulation of Tox, a marker of T cell exhaustion.^37^ Additionally, single-cell T cell receptor (TCR) sequencing revealed more than double the percentage of recurrent TCR clonotypes in the RRV RLI + temozolomide (TMZ) group, indicative of a robust antigen-specific response. The presence of non-virus exposed samples (PBS and PBS + TMZ) in the majority of dominant TCR clusters in combination with the paucity of MLV and pathogen-specific TCR matches or similarities support a tumor-specific reaction rather than one against the viral therapy. These findings, coupled with the increased expression of MHC class I genes in myeloid populations within both the RRV RLI + Vehicle and RRV RLI + TMZ groups, support the hypothesis that TMZ-induced cell death and tumor antigen release in the tumor microenvironment provides additional antigenic material for antigen-presenting cells. These cells, likely further activated by local RLI expression, facilitate cross-presentation to CD8+ T cells. This is further corroborated by CellChat analysis, which demonstrated increased MHC class I–CD8 T cell receptor ligand interactions in the RRV RLI + Vehicle group compared to control, with even greater enhancement observed in the RRV RLI + TMZ group compared to RRV RLI + Vehicle.

This study positions RRV RLI as a potent and clinically translatable viral immunotherapy for GBM, demonstrating robust activation of T cells, NK cells, and enhanced antigen presentation within the TME. Treatment with RRV RLI significantly extends survival in two distinct syngeneic murine GBM models and synergizes effectively with the current standard of care chemotherapy. Furthermore, our TCR-sequencing demonstrating the lack of immunogenicity of the RRV backbone positions RRV as a delivery vehicle for the potent RLI transgene that could potentially be used in repeat treatments in future preclinical studies. Overall, our findings provide compelling preclinical evidence to support a Phase I clinical trial to evaluate the safety and efficacy of RRV RLI in human patients. Moreover, the versatility of this therapeutic approach holds promise for broader application across other cancer types, offering a novel paradigm in the treatment of solid tumors.

## Notes

https://www.ncbi.nlm.nih.gov/geo/query/acc.cgi?acc=GSE278988

### Competing Interest Statement

MKA, NK, AFH, and SC are co-founders and consultants for Kopra Bio, a startup company founded in summer 2024 based on the work presented in this manuscript. NK and SC are employees of 4D Molecular Therapeutics.

### Summary of Updates

We have added the NINDS as a funding source for this work.

### Funding

- National Institute of Neurological Disorders and Stroke: R01NS079697

