## Supplemental Figures for "Non-lytic replicating viral delivery of an IL15 superagonist enhances antitumor immunity and extends survival in glioblastoma"

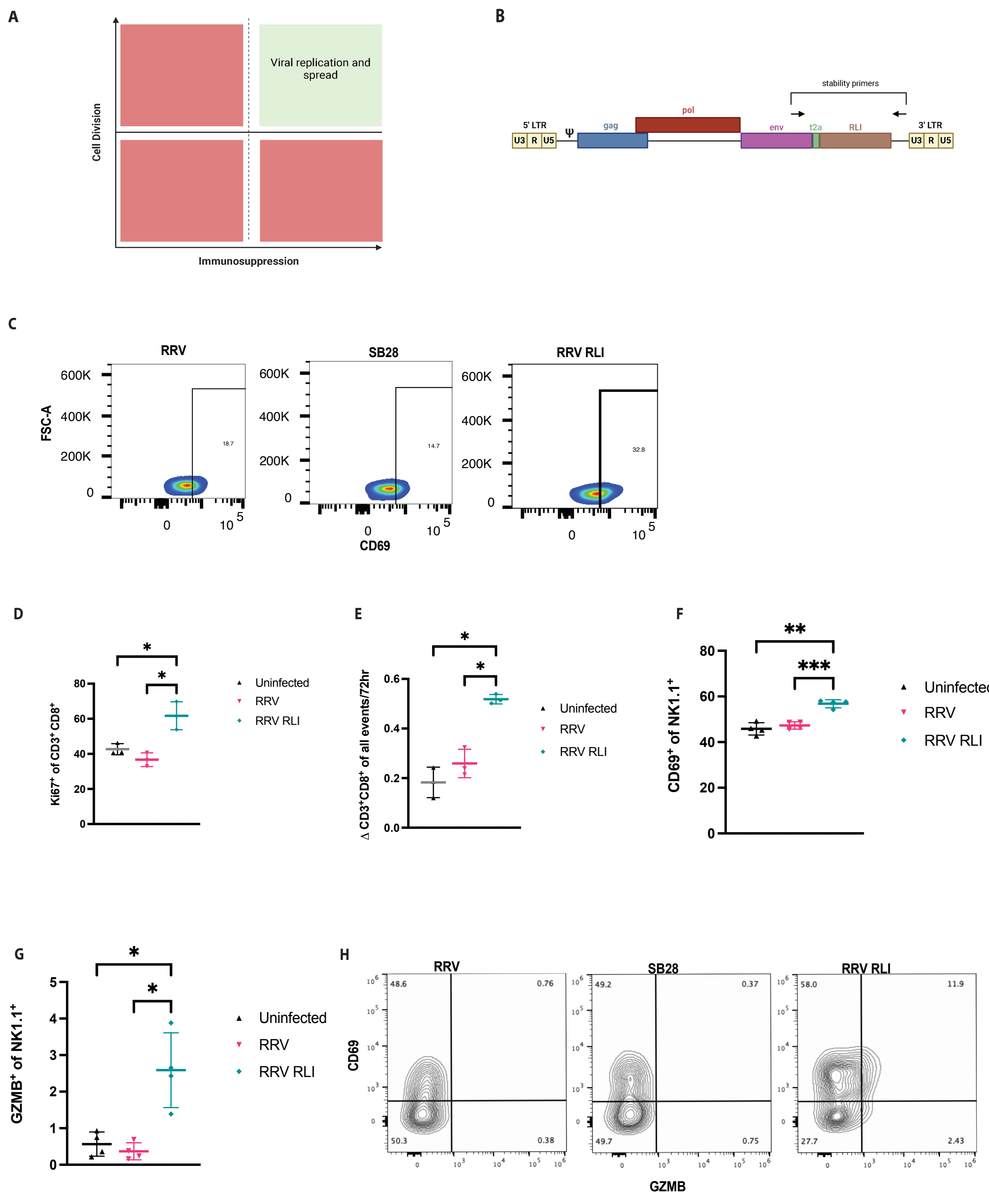

**Fig. S1. RRV RLI rationale and function *in vitro.*** (**A**) RRV RLI relies on cancer hallmarks, cell division and immunosuppression, to form a replication permissive niche. (**B**) Stability primers cross the RLI transgene insertion site, allowing for detection of transgene loss (expected PCR product size: 1018 bp). (**C**) Example gating schematic for CD69 expression in $CD3^{+}CD8^{+}$ T cells in co-culture with infected and uninfected SB28 tumor cells. (**D**) RRV RLI infected SB28 tumor cells increase $Ki67^{+}$ at 72 hours post-addition in co-cultured CD8 T cells relative to RRV and uninfected cells ($p<0.05$, Welch's t-test). (**E**) Co-culture of NK cells with infected SB28 tumor cells demonstrates increased change in $CD8^{+}$ T cell frequency relative to all events at 72 hours when compared to RRV infected and uninfected cells (0.51 vs. 0.26 vs. 0.19, $p<0.05$, Welch's t-test). (**F**) RRV RLI infected SB28 tumor cells increase $CD69^{+}$ single positive cells at 24 hours post-addition in co-cultured NK cells relative to RRV and uninfected cells (56.8% vs. 45.8% vs. 47.2%, $p<0.002$, Welch's t-test). (**G**) RRV RLI infected SB28 tumor cells increase $GZMB^{+}$ single positive cells at 24 hours post-addition in co-cultured NK cells relative to RRV and uninfected cells (2.6% vs. 0.37% vs. 0.57%, $p<0.05$, Welch's t-test). (**H**) Example gating schematic for CD69 and GZMB expression in $CD3^{-}NK1.1^{+}$ cells in co-culture with infected and uninfected SB28 tumor cells.

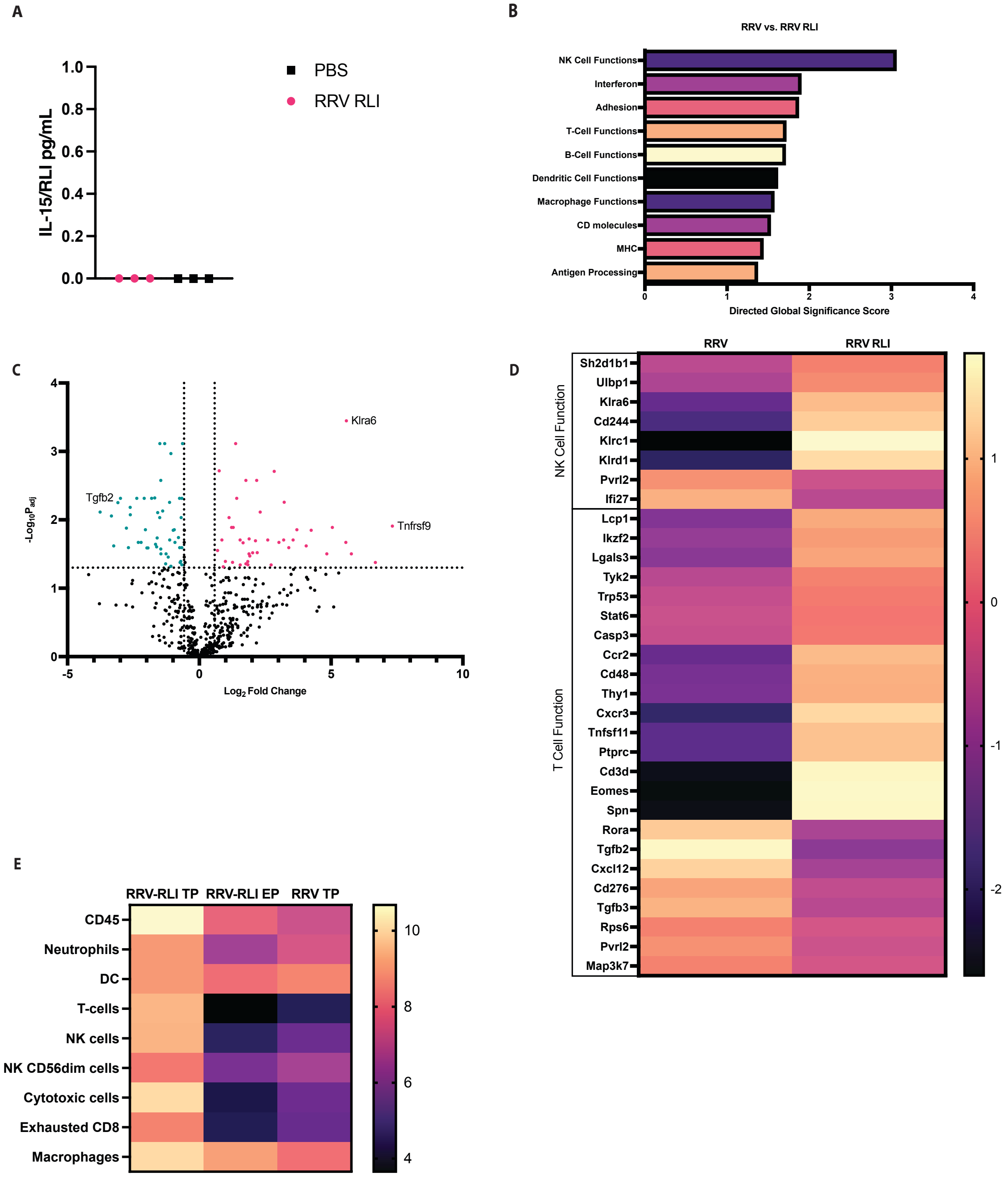
A
IL-15/RLI pg/mL
1.0
0.8
0.6
0.4
0.2
0.0
PBS
RRV RLI
B
RRV vs. RRV RLI
NK Cell Functions
Interferon
Adhesion
T-Cell Functions
B-Cell Functions
Dendritic Cell Functions
Macrophage Functions
CD molecules
MHC
Antigen Processing
0
1
2
3
4
Directed Global Significance Score
C
Klra6
Tgfb2
Tnfrsf9
$-Log_{10}P_{adj}$
$Log_2$ Fold Change
-5
0
5
10
D
RRV
RRV RLI
NK Cell Function
Sh2d1b1
Ulbp1
Klra6
Cd244
Klrc1
Klrd1
Pvrl2
Ifi27
T Cell Function
Lcp1
Ikzf2
Lgals3
Tyk2
Trp53
Stat6
Casp3
Ccr2
Cd48
Thy1
Cxcr3
Tnfsf11
Ptprc
Cd3d
Eomes
Spn
Rora
Tgfb2
Cxcl12
Cd276
Tgfb3
Rps6
Pvrl2
Map3k7
1
0
-1
-2
E
RRV-RLI TP
RRV-RLI EP
RRV TP
CD45
Neutrophils
DC
T-cells
NK cells
NK CD56dim cells
Cytotoxic cells
Exhausted CD8
Macrophages
10
8
6
4

**Fig S2. Local RRV RLI therapy modulates immune response by affecting T cells, NK cells, and antigen presentation.** (**A**) No systemic IL-15/RLI is present in the blood of treated SB28 tumor bearing mice at day 14 after tumor implantation (day 10 after virus injection). (**B**) Directed global significance scores of the top 10 upregulated gene pathways in RRV RLI treatment vs. RRV treatment. (**C**) Volcano plot showing differentially expressed genes between RRV RLI and RRV treatment groups at day 14 post tumor implantation timepoint. (**D**) Heat map detailing differential gene expression for genes related to T cell function and NK cell function between RRV RLI and PBS treatment groups. (**E**) Heat map demonstrating calculated immune infiltration scores between RRV RLI at day 14 post tumor implantation timepoint (RRV RLI TP), RRV RLI at endpoint (RRV RLI EP), and RRV at day 14 post tumor implantation timepoint (RRV TP).

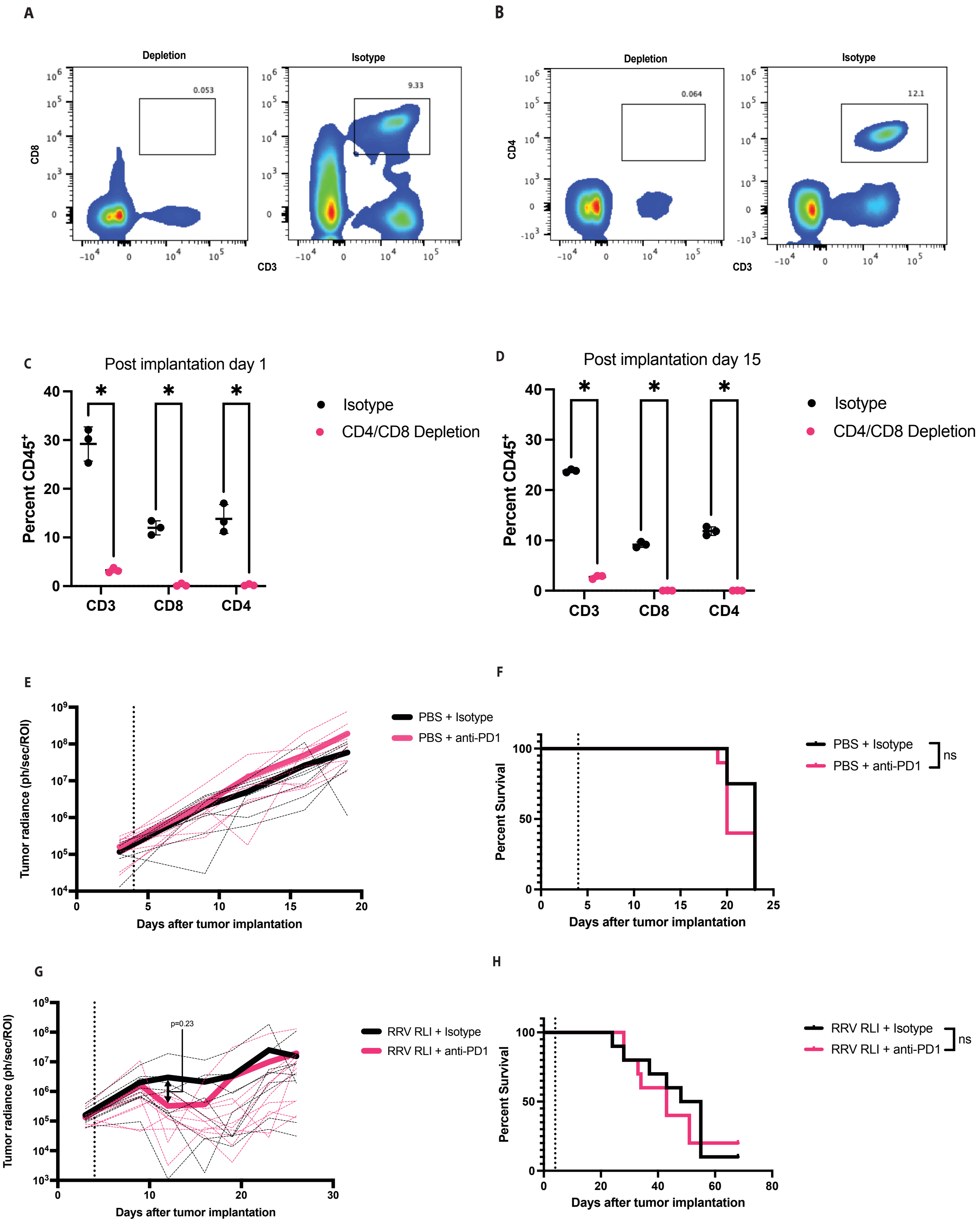
A
Depletion
Isotype
CD8
CD3
0.053
9.33
B
Depletion
Isotype
CD4
CD3
0.064
12.1
C
Post implantation day 1
Percent CD45+
CD3
CD8
CD4
Isotype
CD4/CD8 Depletion
D
Post implantation day 15
Percent CD45+
CD3
CD8
CD4
Isotype
CD4/CD8 Depletion
E
Tumor radiance (ph/sec/ROI)
Days after tumor implantation
PBS + Isotype
PBS + anti-PD1
F
Percent Survival
Days after tumor implantation
PBS + Isotype
PBS + anti-PD1
ns
G
Tumor radiance (ph/sec/ROI)
p=0.23
Days after tumor implantation
RRV RLI + Isotype
RRV RLI + anti-PD1
H
Percent Survival
Days after tumor implantation
RRV RLI + Isotype
RRV RLI + anti-PD1
ns

**Fig S3. Validation of immune cell depletion and demonstration that RRV RLI combination with anti-PD1 treatment does not alter therapeutic efficacy.**
(**A**) Example flow cytometry gaiting for the validation of CD4 and CD8 depletion studies. (**B**) Flow cytometric analysis of anti-CD4 and anti-CD8 antibody-treated SB28 tumor bearing mice reveals successful depletion of $CD3^+$ (3.2% vs. 29.2%, $p<0.01$, Welch's t-test), $CD8^+$ (0.2% vs. 12.0%, $p<0.01$, Welch's t-test), and $CD4^+$ T cells (0.23% vs. 13.8%, $p<0.02$, Welch's t-test), relative to isotype control at day 1 post tumor implantation. (**C**) Similar results were seen for $CD3^+$ (2.7% vs. 23.8%, $p<0.001$, Welch's t-test), $CD8^+$ (0.04% vs. 9.1%, $p<0.01$, Welch's t-test), and $CD4^+$ T cells (0.03% vs. 11.8%, $p<0.02$, Welch's t-test), relative to isotype control at day 15 post tumor implantation. (**D**) In PBS-treated mice bearing SB28 tumors, anti-PD1 therapy did not reduce tumor growth on bioluminescent imaging compared to isotype-treated controls. (**E**) In PBS-treated mice bearing SB28 tumors, anti-PD1 therapy did not extend survival compared to isotype-treated controls (median survival 20 days vs. 23 days; $p = 0.13$, Log-Rank Mantel-Cox test). (**F**) In RRV RLI-treated mice bearing SB28 tumors, anti-PD1 therapy did not reduce tumor growth on bioluminescent imaging compared to isotype-treated controls ($p=0.23$, day 12 post tumor implantation, Welch's t-test) (**E**) In RRV RLI-treated mice bearing SB28 tumors, anti-PD1 therapy did not extend survival compared to isotype-treated controls (median survival 43 days vs. 51 days; $p = 0.70$, Log-Rank Mantel-Cox test).

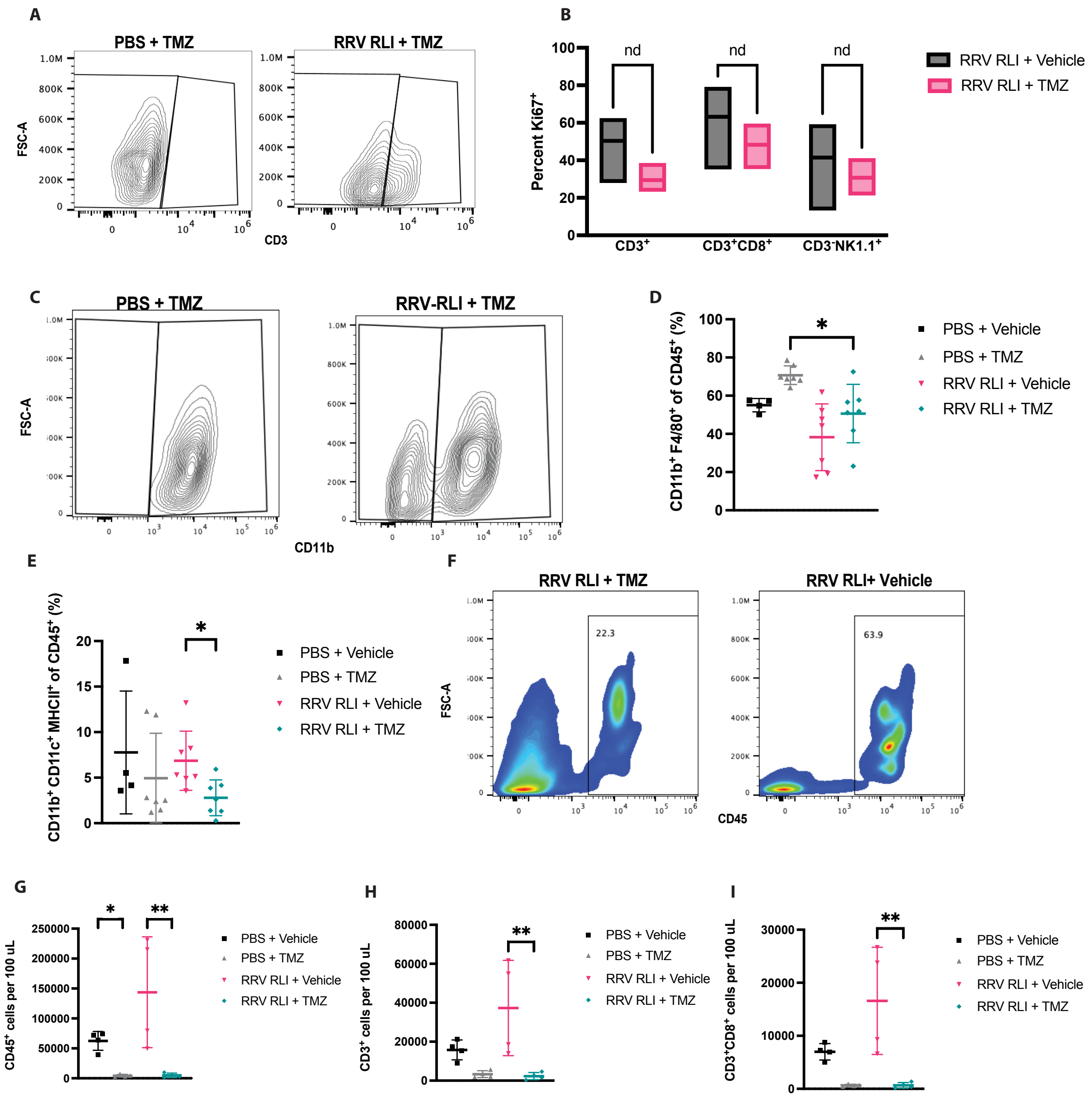
A
PBS + TMZ
RRV RLI + TMZ
FSC-A
CD3
B
Percent Ki67+
nd
RRV RLI + Vehicle
RRV RLI + TMZ
CD3+
CD3+CD8+
CD3-NK1.1+
C
PBS + TMZ
RRV-RLI + TMZ
CD11b
D
CD11b+ F4/80+ of CD45+ (%)
PBS + Vehicle
PBS + TMZ
RRV RLI + Vehicle
RRV RLI + TMZ
E
CD11b+ CD11c+ MHCII+ of CD45+ (%)
F
RRV RLI + TMZ
RRV RLI+ Vehicle
22.3
63.9
CD45
G
CD45+ cells per 100 uL
H
CD3+ cells per 100 uL
I
CD3+CD8+ cells per 100 uL

**Fig S4. Additional immune phenotyping reveals systemic TMZ associated myelosuppression.** (**A**) Example gating strategy for $CD3^+$ cells (already gated on live $CD45^+CD11b^-$). (**B**) Both RRV RLI + Vehicle and RRV RLI + TMZ result in significant $Ki67^+$ $CD3^+$, $CD3^+CD8^+$, and $CD3^-NK1.1^+$ populations with no difference between the groups. (**C**) Example gating strategy for $CD11b^+$ cells (already gated on live $CD45^+$). (**D**) Percent tumor infiltrating $CD11b^+F4/80^+$ cells with a small reduction in RRV RLI + TMZ vs. PBS + TMZ ($p<0.05$, Uncorrected Dunn's test). (**E**) Percent tumor infiltrating $CD11b^+CD11c^+MHCII^+$ cells with a small reduction in RRV RLI + TMZ vs. RRV RLI + Vehicle ($p<0.05$, Uncorrected Dunn's test). (**F**) Example gating strategy for $CD45^+$ cells in SB28 tumor bearing mice blood samples (already gated on live single cells). (**G**) $CD45^+$ counts per 100 uL in SB28 tumor bearing mice blood samples with reductions in total cells per 100 uL in PBS + TMZ vs. PBS + Vehicle ($p<0.05$, Uncorrected Dunn's test) and RRV RLI + TMZ vs. RRV RLI + Vehicle ($p<0.001$, Uncorrected Dunn's test) (**H**) $CD3^+$ counts per 100 uL in SB28 tumor bearing mice blood samples with a reduction in total cells per 100 uL in RRV RLI + TMZ vs. RRV RLI + Vehicle ($p<0.001$, Uncorrected Dunn's test) (**I**) $CD3^+CD8^+$ counts per 100 uL in SB28 tumor bearing mice blood samples with a reduction in total cells per 100 uL in RRV RLI + TMZ vs. RRV RLI + Vehicle ($p<0.001$, Uncorrected Dunn's test).

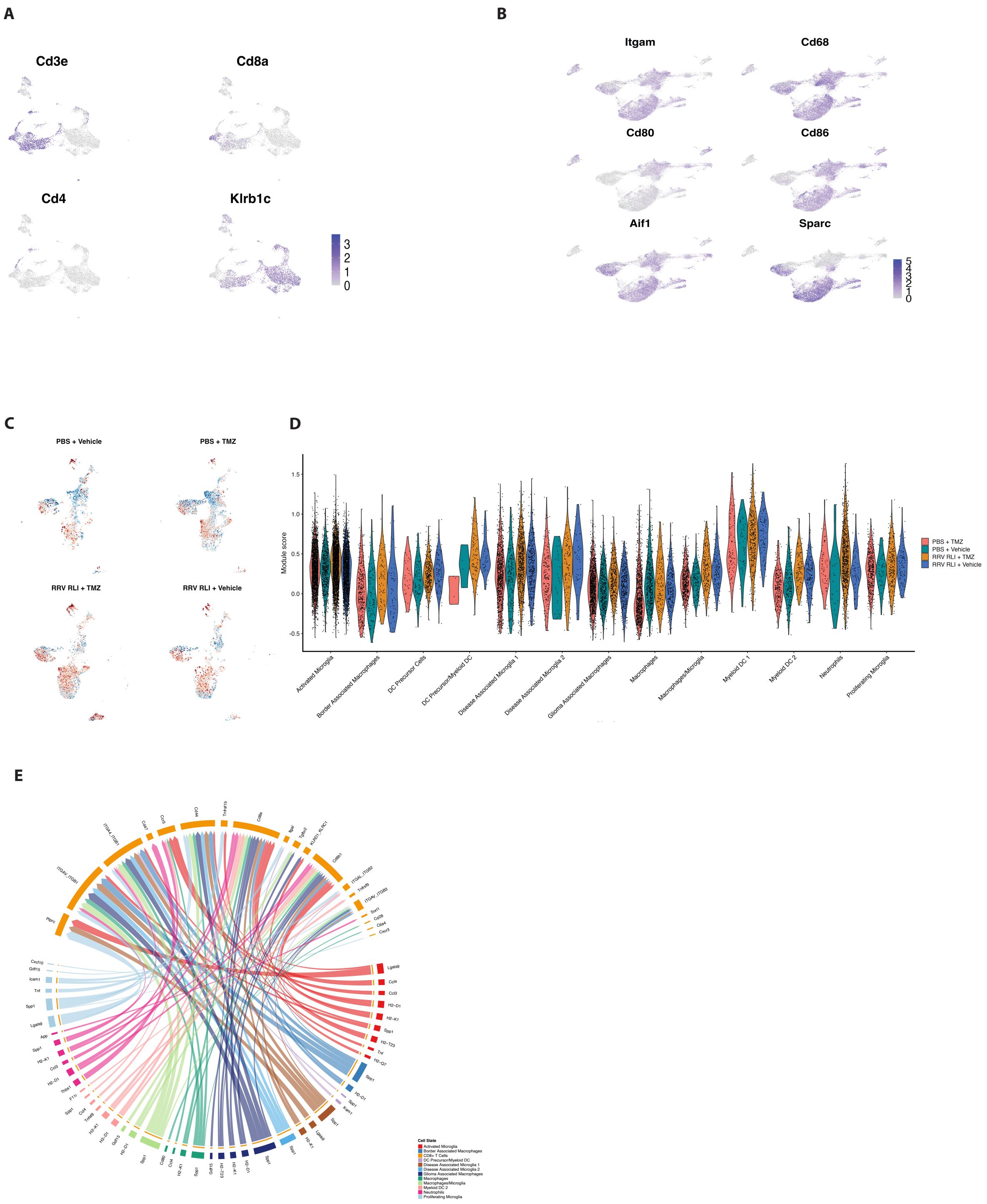
A
Cd3e
Cd8a
Cd4
Klrb1c
3
2
1
0
B
Itgam
Cd68
Cd80
Cd86
Aif1
Sparc
C
PBS + Vehicle
PBS + TMZ
RRV RLI + TMZ
RRV RLI + Vehicle
D
Module score
1.5
1.0
0.5
0.0
-0.5
Activated Microglia
Border Associated Macrophages
DC Precursor Cells
DC Precursor/Myeloid DC
Disease Associated Microglia 1
Disease Associated Microglia 2
Glioma Associated Macrophages
Macrophages
Macrophages/Microglia
Myeloid DC 1
Myeloid DC 2
Neutrophils
Proliferating Microglia
PBS + TMZ
PBS + Vehicle
RRV RLI + TMZ
RRV RLI + Vehicle
E
Cell State
Activated Microglia
Border Associated Macrophages
CD8+ T Cells
DC Precursor/Myeloid DC
Disease Associated Microglia 1
Disease Associated Microglia 2
Glioma Associated Macrophages
Macrophages
Macrophages/Microglia
Myeloid DC 2
Neutrophils
Proliferating Microglia

**Fig S5. Single-cell RNA sequencing identifies enhanced effector cell activation and antigen presentation driving efficacy in combined RRV RLI and TMZ therapy.** (**A**) UMAP of all sequenced T and NK cells demonstrating key marker gene expression. (**B**) UMAP of all sequenced myeloid cells demonstrating key marker gene expression. (**C**) UMAP of all sequenced myeloid cells demonstrating changes in MHC class I gene expression score between samples. (**D**) Violin plot of MHC class I gene expression score in myeloid subpopulations between samples. (**E**) Cell chat analysis of incoming signaling in RRV RLI + TMZ vs. RRV RLI + Vehicle.

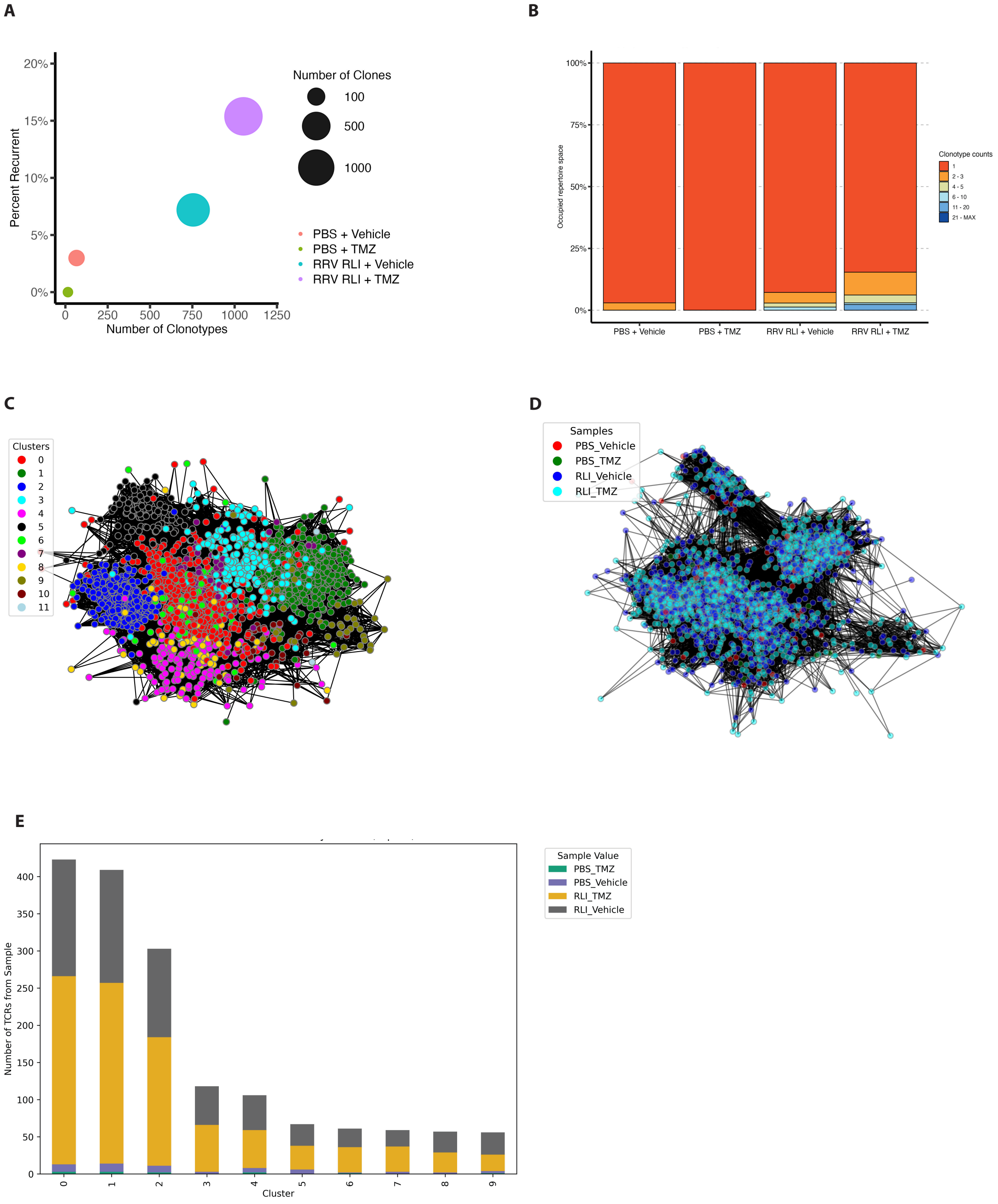
A
B
C
D
E
Number of Clones
100
500
1000
PBS + Vehicle
PBS + TMZ
RRV RLI + Vehicle
RRV RLI + TMZ
Percent Recurrent
Number of Clonotypes
20%
15%
10%
5%
0%
0
250
500
750
1000
1250
Occupied repertoire space
Clonotype counts
1
2 - 3
4 - 5
6 - 10
11 - 20
21 - MAX
100%
75%
50%
25%
0%
PBS + Vehicle
PBS + TMZ
RRV RLI + Vehicle
RRV RLI + TMZ
Clusters
0
1
2
3
4
5
6
7
8
9
10
11
Samples
PBS_Vehicle
PBS_TMZ
RLI_Vehicle
RLI_TMZ
Sample Value
PBS_TMZ
PBS_Vehicle
RLI_TMZ
RLI_Vehicle
Number of TCRs from Sample
Cluster
400
350
300
250
200
150
100
50
0
0
1
2
3
4
5
6
7
8
9

**Fig S6. Single-cell TCR sequencing suggests tumor-specific T cell clonal expansion induced by RRV RLI and temozolomide therapy.** (**A**) X-Y graph showing increased recurrent T cell clones in RRV RLI + TMZ vs. RRV RLI + Vehicle. (**B**) Visualization of clone frequency occupancy by clone rank (**C**) Beta chain clustering of TCRs from all samples within 80 distance units with specific clusters labeled. (**D**) Beta chain clustering of TCRs from all samples within 80 distance units with labeling of sample specific TCRs. (**E**) Stacked bar graph demonstrating sample identification across the top 9 beta chain clusters.
